# An integrated transcriptomic and epigenomic atlas of mouse primary motor cortex cell types

**DOI:** 10.1101/2020.02.29.970558

**Authors:** Zizhen Yao, Hanqing Liu, Fangming Xie, Stephan Fischer, A. Sina Booeshaghi, Ricky S Adkins, Andrew I. Aldridge, Seth A. Ament, Antonio Pinto-Duarte, Anna Bartlett, M. Margarita Behrens, Koen Van den Berge, Darren Bertagnolli, Tommaso Biancalani, Héctor Corrada Bravo, Tamara Casper, Carlo Colantuoni, Heather Creasy, Kirsten Crichton, Megan Crow, Nick Dee, Elizabeth L Dougherty, Wayne I. Doyle, Sandrine Dudoit, Rongxin Fang, Victor Felix, Olivia Fong, Michelle Giglio, Jeff Goldy, Mike Hawrylycz, Hector Roux de Bézieux, Brian R. Herb, Ronna Hertzano, Xiaomeng Hou, Qiwen Hu, Jonathan Crabtree, Jayaram Kancherla, Matthew Kroll, Kanan Lathia, Yang Eric Li, Jacinta D. Lucero, Chongyuan Luo, Anup Mahurkar, Delissa McMillen, Naeem Nadaf, Joseph R. Nery, Sheng-Yong Niu, Joshua Orvis, Julia K. Osteen, Thanh Pham, Olivier Poirion, Sebastian Preissl, Elizabeth Purdom, Christine Rimorin, Davide Risso, Angeline C. Rivkin, Kimberly Smith, Kelly Street, Josef Sulc, Thuc Nghi Nguyen, Michael Tieu, Amy Torkelson, Herman Tung, Eeshit Dhaval Vaishnav, Valentine Svensson, Charles R. Vanderburg, Vasilis Ntranos, Cindy van Velthoven, Xinxin Wang, Owen R. White, Z. Josh Huang, Peter V. Kharchenko, Lior Pachter, John Ngai, Aviv Regev, Bosiljka Tasic, Joshua D. Welch, Jesse Gillis, Evan Z. Macosko, Bing Ren, Joseph R. Ecker, Hongkui Zeng, Eran A. Mukamel

## Abstract

Single cell transcriptomics has transformed the characterization of brain cell identity by providing quantitative molecular signatures for large, unbiased samples of brain cell populations. With the proliferation of taxonomies based on individual datasets, a major challenge is to integrate and validate results toward defining biologically meaningful cell types. We used a battery of single-cell transcriptome and epigenome measurements generated by the BRAIN Initiative Cell Census Network (BICCN) to comprehensively assess the molecular signatures of cell types in the mouse primary motor cortex (MOp). We further developed computational and statistical methods to integrate these multimodal data and quantitatively validate the reproducibility of the cell types. The reference atlas, based on more than 600,000 high quality single-cell or -nucleus samples assayed by six molecular modalities, is a comprehensive molecular account of the diverse neuronal and non-neuronal cell types in MOp. Collectively, our study indicates that the mouse primary motor cortex contains over 55 neuronal cell types that are highly replicable across analysis methods, sequencing technologies, and modalities. We find many concordant multimodal markers for each cell type, as well as thousands of genes and gene regulatory elements with discrepant transcriptomic and epigenomic signatures. These data highlight the complex molecular regulation of brain cell types and will directly enable design of reagents to target specific MOp cell types for functional analysis.

## Introduction

Neural circuits are characterized by extraordinary diversity of their cellular components^1, 2^. Single-cell molecular assays, especially transcriptomic measurements by RNA-Seq, have accelerated the discovery and characterization of cell types across brain regions and in diverse species. Recent advances include single-cell transcriptome datasets with >10^5^ individual cells, identifying hundreds of neuronal and non-neuronal cell types across the mouse nervous system^3–5^. As the number of profiled cells grows into the millions, a key question is whether these data will converge toward a comprehensive and coherent taxonomy of cell types with broad utility for organizing knowledge of brain cells and their function. Data from different modalities, including transcriptomic and epigenomic data, must be cross-referenced and integrated to establish robust and consistent cell type classifications. Although a comprehensive atlas should incorporate anatomical and physiological information, the high throughput of single cell sequencing assays makes integration of molecular data a particularly urgent challenge. A rigorous and reproducible consensus molecular atlas of brain cell types would drive progress across modalities, including obtaining functional information.

Single cell sequencing technologies can measure multiple molecular signatures of cell identity. The core molecular identity of a cell is largely established during development and maintained by a combination of gene regulatory proteins, such as transcription factors, and epigenetic marks, such as open chromatin and DNA methylation^6,7^. The expression of specific cell fate-determining proteins promotes stable, covalent modifications of chromatin and DNA, while epigenetic marks in turn shape and maintain cell type-specific gene expression. Transcription and epigenetic modifications, acting on timescales from minutes to decades, mutually reinforce each other and establish attractors in cellular state space corresponding to cell types^8–10^. Neurons express a range of cell type marker genes and gene modules that shape their mode of synaptic communication^11^, but cell state and gene expression can also vary due to circadian rhythms^12^ and neural activity^13^. Neuronal DNA methylation is reconfigured during an extended postnatal period^14,15^, leading to highly cell type-specific patterns of CG and non-CG methylation in mature neurons^16,17^. In addition, the physical configuration of DNA, especially the locations of open chromatin regions, correlates with cell type-specific gene regulation and provides rich cell identity information^18^. Several technologies now enable measurement of these molecular signatures in thousands to hundreds of thousands of individual cells or nuclei, generating large-scale datasets that are both wide (many features) and deep (many cells). Here, we integrate such cell type signatures to achieve a reference taxonomy for one brain region, the adult mouse primary motor cortex, using a combination of single cell and single nucleus transcriptomes, DNA methylomes and open chromatin datasets. This atlas represents a first step toward the goal of the BICCN to generate a comprehensive cell-type atlas comprising all regions of the mouse brain^19^.

## Results

### A multimodal approach to molecular atlasing of mouse primary motor cortex (MOp)

We aimed to comprehensively identify and characterize the molecular identity of all cell types in the adult mouse primary motor cortex (Fig 1a,b). To achieve this, we formed a collaborative network within the framework of the BICCN to coordinate collection of single-cell and single-nucleus samples followed by sequencing. We brought together 9 separate datasets, including 7 single-cell or single-nucleus transcriptome datasets (single-cell and single-nucleus RNA-seq using 10x v2, v3 and SMART-Seq; n=732,779 cells), one single-nucleus DNA methylation dataset (snmC-Seq2, n=9,941) and one single-nucleus open chromatin dataset (snATAC-Seq, n=135,665) (Supplementary Table 1). These datasets span a range of technologies with complementary strengths, including different number of cells assayed, depth of sequence coverage per cell, and biological features assessed (Fig. 1c,d). The datasets we produced reflect the inherent tradeoff in single cell sequencing assays between number of sequenced molecules per cell, which corresponds to sequencing depth, and the total number of cells that can be assayed for a fixed total cost. At one end of this spectrum, our datasets include a large set of single-nucleus transcriptomes from over 175,000 cells (using the 10x Chromium 3’ version 3 platform). By contrast, our full-length transcript sequencing using SMART-Seq v4 captured a greater number of genes per cell, but covered fewer cells (∼6,500 per dataset). Single-nucleus DNA methylation data provided deep coverage of the epigenome per cell for a modest number of cells^16,20^ (∼10,000), whereas snATAC-Seq data scaled to over 100,000 cells but sampled fewer DNA fragments for individual cells^18^.

**Figure 1:**
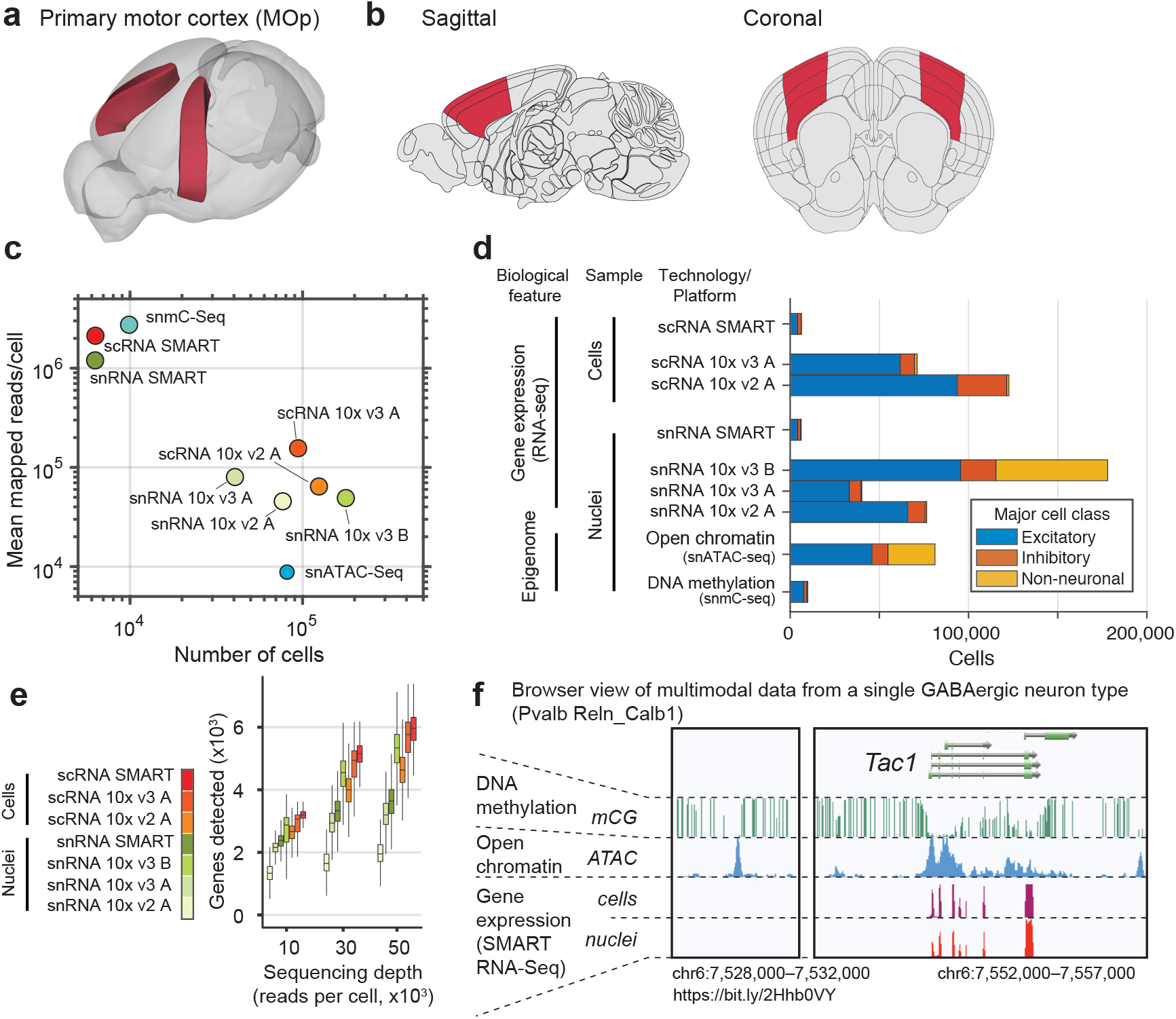
A multimodal molecular cell type atlas of mouse primary motor cortex (MOp). **a,** Anatomical location of mouse MOp in the Allen Mouse Brain Common Coordinate Framework (CCFv3). **b,** Representative sagittal and coronal sections and dissected MOp region. **c,** Number of cells and number of sequencing reads per cell in each of 9 single-cell transcriptome and epigenome datasets. **d,** Number of cells in each of the major cell classes {glutamatergic excitatory, GABAergic inhibitory neurons, non-neurons) of each dataset (excluding snRNA 10x v2 A). Differences in cell type sampling strategy affect the relative number of neurons and non-neuronal cells. **e,** Number of genes detected per cell or nucleus for transcriptomic data following a down-sampling analysis of sequencing depths. **f,** Example genome browser tracks for the *Tac1* gene comparing three data modalities for one cell type.

Subsampling analysis of RNA-Seq datasets (Fig. 1e) shows that in general, scRNA-Seq (both SMART and 10x) detects more genes per cell than snRNA-Seq, and the 10x v3 platform performs substantially better than 10x v2. An interesting exception is that the number of genes detected per cell in the snRNA-Seq 10x v3 B dataset, using an improved nucleus isolation protocol^21^, is significantly higher than those of all other snRNA-Seq datasets, and is comparable to that of the scRNA-Seq 10x v3 dataset.

To illustrate the correspondence among different technologies, sampling strategies, and data modalities, we highlight the Tachykinin-1 gene locus (*Tac1,* Fig. 1f; <u>browser</u>). This gene is a specific marker of a subset of medial ganglionic eminence (MGE)-derived inhibitory GABAergic interneurons. Our data confirm the expression of *Tac1* mRNA in a cluster of parvalbumin-expressing neurons, with RNA transcripts captured in both single-cell and single-nucleus preparations. We further observed accessible chromatin and low DNA methylation at CG sites within the body of the *Tac1* gene, and at an intergenic location ∼20 kb upstream of the transcription start site. Our study takes advantage of such multimodal, multi-scale, cell type-specific molecular signatures to build a comprehensive transcriptomic and epigenomic atlas of the mouse MOp.

We have created several web resources to enable interactive data access, interactive exploration, visualization, and analysis (Extended Data Fig. 1). Raw sequence data are available at the Neuroscience Multi-omics Archive (nemoarchive.org). A suite of web-based tools for visualization and analysis of the integrated transcriptomic and epigenomic data are available at NeMO Analytics (nemoanalytics.org) and the brainome portal (brainome.ucsd.edu/BICCN_MOp). These portals allow users to visualize integrated multi-omic data across experiments and species side-by-side via genome and cell browsers, perform cluster comparison, identify marker genes.

### A consensus transcriptomic atlas based on multiple single cell and nucleus RNA-Seq datasets

To establish a transcriptomic reference atlas of mouse MOp and to directly compare with existing cell taxonomies, we jointly analyzed 7 single-cell (sc) and single-nucleus (sn) RNA-Seq datasets. The datasets were mutually consistent, with strongly correlated expression of cell type marker genes (Extended Data Fig. 2a,c,d) despite differences in the sensitivity to genes with low expression (Extended Data Fig. 2b). Computational data integration using *scrattch.hicat* (Methods), which adjusts for systematic differences between datasets due to technical differences or uncontrolled batch effects, enabled clustering and identification of 116 cell types (Fig. 2a, Extended Data Fig. 2c, Supplementary Tables 2-4). Importantly, cells and nuclei assayed by each of the technologies and in each batch grouped primarily by cell type and not by dataset (Fig. 2b). Residual dataset-related differences, including systematic differences between nuclear and cellular RNA-Seq assays, could be observed in some clusters as a gradient of transcriptomes from different datasets. We performed hierarchical clustering of the cell types based on average gene expression for each cell type to uncover the the relationships among types within each major cell class: GABAergic inhibitory neurons (n=59 types), glutamatergic excitatory neurons (n=31) and non-neurons (n=26) (Fig. 2c). Six of the transcriptomic datasets used cell sorting strategies to enrich neurons relative to non-neuronal cells, while the largest dataset (snRNA 10x v3 B) represents an unbiased sampling of both neuronal and non-neuronal cells. Despite these differences, the relative frequency of cell types was highly consistent across datasets after normalizing for the total sample of each major class (Fig. 2d). 86 out of 116 cell types were present across all of the RNA-Seq datasets, while the rest are either non-neuronal types that were under-sampled in many datasets, or extremely rare types (< 0.01% of all cells).

**Figure 2:**
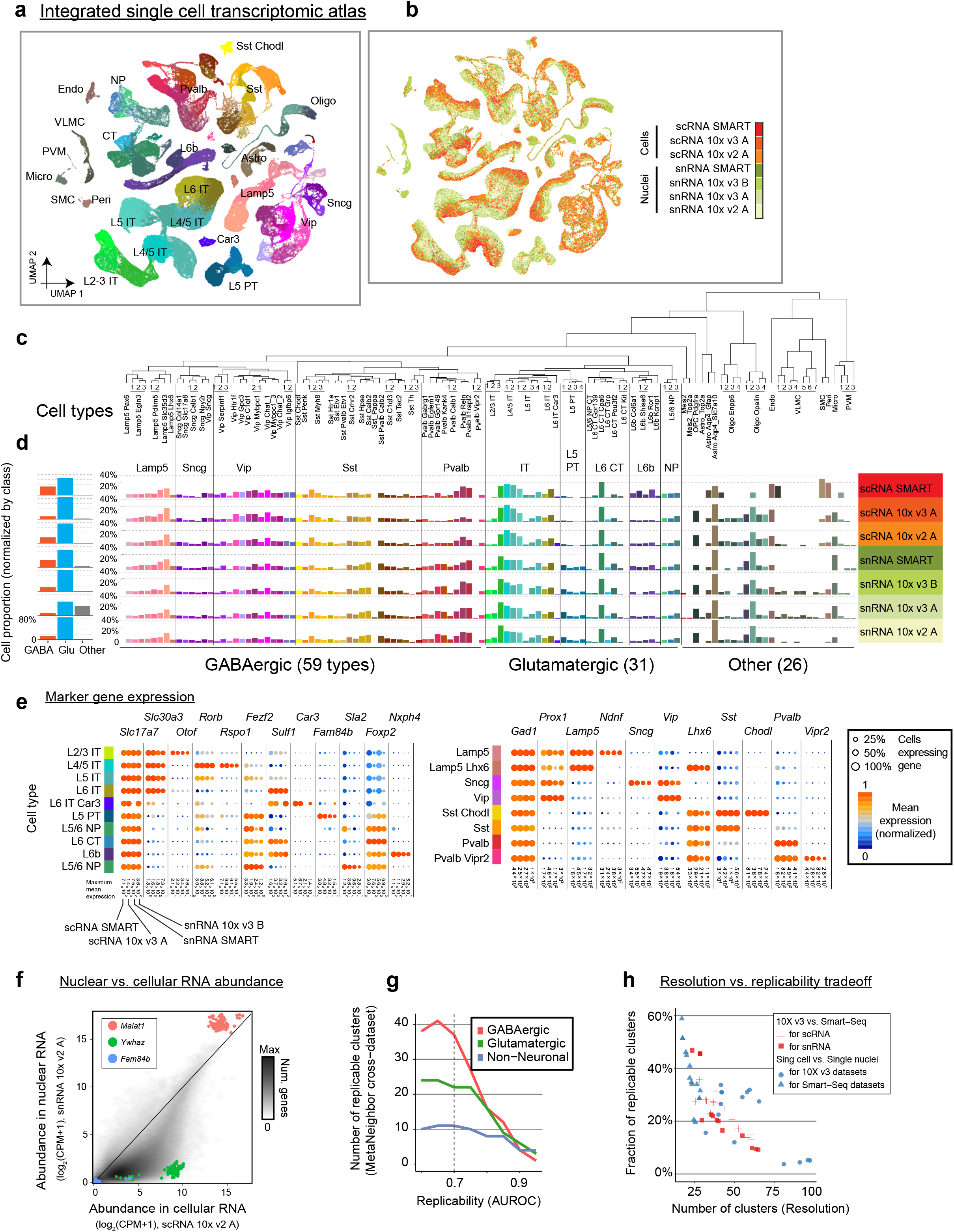
Multi-platform integrated transcriptomic taxonomy of MOp cell types. **a,b** Two-dimensional projection (UMAP) of cells and nuclei based on integrated analysis of seven datasets using Seurat, followed by cluster analysis. Individual cells and nuclei are colored by cell type (**a**), or by data platform (**b**). Non-neuronal cell types are depleted in all datasets except snRNA 10x v3 B due to the sampling strategy, which enriched neurons. **c,** Dendrogram showing hierarchical relationship among the consensus transcriptomic cell types. **d**, Proportion of cells of each type per dataset, normalized within major classes. **e,** Expression of marker genes for excitatory and inhibitory cell classes, across four platforms. **f,** Differential enrichment of transcripts in single cells (x-axis) vs. single nuclei (y-axis). Non-coding RNAs such as *Malat1* are enriched in nuclei. **g,** Number of clusters replicable across at least two of the seven sc/snRNA-seq datasets as a function of minimal MetaNeighbor score. **h,** Trade-off between number of clusters and replicability (fraction of clusters with minimal MetaNeighbor replicability score). Lamp5/Sncg/Vip/Sst/Pvalb - Major inhibitory neuron subclasses; L2-6 - layers; IT - Intratelencephalic; PT - Pyramidal tract; CT - Corticothalamic; NP - Near-projecting; Astro - Astrocytes; OPC - Oligodendrocyte precursor; Oligo - Oligodendrocytes; Micro - Microglial cells; SMC - Smooth muscle cells; VLMC - Vascular lepotomeningeal cells; Peri - Pericyte; PVM - Perivascular macrophage; Endo – Endothelial

The transcriptomic MOp cell taxonomy is a data-driven resource, with objective and quantitative signatures of each cell type that can be used to compare with existing and forthcoming datasets (Supplementary Table 5). To facilitate the use of these cell types by investigators, we adopted a nomenclature that incorporates multiple anatomic and molecular identifiers. For example, we identified four clusters of excitatory neurons (expressing *Slc17a7* encoding vesicular glutamate transporter Vglut1) that express markers of deep layers (*Fezf2, Foxp2*) as well as *Fam84b*, a unique marker of extra-telencephalic projecting neurons^22^ (pyramidal tract, PT) (Fig. 2e). These neurons were therefore labeled “L5 PT 1-4”. For GABAergic neurons, our nomenclature divides cells into 5 major subclasses based on marker genes (*Lamp5*, *Sncg*, *Vip*, *Sst*, *Pvalb*), with finer clusters identified by secondary markers (e.g. *Sst*, *Myh8*). To track each of these clusters and uniquely associate them with the underlying molecular data, we provide accession IDs compatible with a proposed cell type nomenclature and a full list of the top marker genes for every pair of cell types (Supplementary Table 3,6)^23^.

To facilitate annotation of MOp cell types and comparison with other cortical regions, we assigned each single cell or nucleus to the best matching cell type in a large dataset (n=23,822 SMART-Seq cells) of mouse anterolateral motor cortex (ALM) and primary visual cortex (VISp) neurons (Extended Data Fig. 3a)^5^. We found one-to-one matches between most of the 116 MOp cell types and the 102 previously defined cortical cell types in ALM. In particular, we found 4 types of Layer 5 pyramidal tract (L5 PT) neurons, which correspond with 3 previously described^24^ deep layer excitatory neuron types with distinct subcortical projection patterns to thalamus and medulla (Extended Data Fig. 3b,c). These types, which were associated with distinct roles in movement planning and initiation, were distinguished by robust patterns of differential gene expression in each of the transcriptomic datasets (Extended Data Fig. 4).

The motor cortex is traditionally considered to lack a discernible layer 4 based on the absence of a clear cytoarchitectonic signature^25^. However, recent anatomical studies have identified a population of pyramidal cells located between layers 3 and 5, with hallmarks of L4 neurons including thalamic input and outputs to L4 and L2/3^26^. We identified two clusters, containing over 99,000 cells, which express a combination of markers usually associated with L4^27^, including *Cux2*, *Rspo1* and *Rorb* (both clusters), and those associated with L5, *e.g.*, *Fezf2* (one cluster) (Fig. 2e, Extended Data Fig. 5). We therefore labeled these clusters L4/5.

By collecting both scRNA-Seq and snRNA-Seq data, using multiple platforms and with high sampling depth, we could directly compare the nuclear and cytoplasmic transcriptomes of MOp cells. A comparison of sc- and snRNA-Seq on a smaller scale in mouse visual cortex (VISp) showed that both modalities can provide comparable clustering resolution^28^, consistent with our analyses of individual datasets (Extended Data Fig. 2a,b). L4/5 cells in MOp have a larger proportion of nuclear transcripts than L5 IT and L5 PT cells, consistent with previous observations in VISp^28^ (Extended Data Fig. 2f). We further examined whether individual genes are enriched in the nuclear or cytoplasmic RNA fraction across MOp cell types, finding that scRNA and snRNA protocols reveal differences in mRNA localization (Fig. 2f, Extended Data Fig. 2e-g). For example, the long non-coding RNA *Malat1* was enriched in snRNA-Seq, consistent with its known nuclear localization^29^. By contrast, mRNA of the protein-coding gene *Ywhaz* was strongly depleted from the nucleus. This result complements a recent observation of specific localization of *Ywhaz* mRNA in the somata, but not the dendrites, of hippocampal neurons^30^.

### Statistical reproducibility of transcriptomic clusters across datasets

Single-cell sequencing has enabled a proliferation of transcriptomic, epigenomic and multimodal studies of brain cell types. To make progress, these separate datasets should be compared and integrated using objective and meaningful biological and statistical criteria. Our mouse MOp datasets represent the most comprehensive collection of single-cell datasets from a single region to date, providing an unprecedented opportunity to investigate the statistical reproducibility and robustness of cell taxonomies across a broad range of technical parameters and data modalities. We applied MetaNeighbor to assess the cross-dataset replicability of clusters defined separately using each of the seven transcriptomic datasets (Supplementary Table 4)^31^. This analysis tests whether cell types defined using one dataset can be predicted by using the closest matching (nearest neighbor) cells in other datasets, together with the independent cluster results for the cells in the other datasets. We found 70 clusters with a high statistical replicability score (AUROC > 0.7 across at least two out of seven datasets, Fig. 2g), including 37 GABAergic neurons, 22 glutamatergic, and 11 non-neuronal cell types. Most of the clusters had reciprocal best matches across all datasets investigated or were missing in only one dataset (Extended Data Fig. 7a).

MetaNeighbor analysis further allowed us to examine the consistency of different computational clustering procedures. We ran three widely used single-cell analysis packages ^32–34^ to generate a fine-grained clustering of each dataset. These cluster analyses were not optimized or manually curated; instead, we used “off-the-shelf” computational procedures to test the robustness of the results from a relatively straightforward and automated analysis. These clusters are thus expected to be less biologically meaningful and robust compared with more customized procedures, such as our reference clustering that incorporates analysis of differential expression to validate the biological reality of cell types. Using the three off-the-shelf cluster analyses, we created a sequence of increasingly coarse-grained clusterings by iteratively merging pairs of clusters chosen to maximize the consistency across computational methods (ARI-merging; see Methods). Finally, at each level of resolution we used MetaNeighbor to calculate the number of clusters which were highly replicable (AUROC>0.7) across datasets. The result of this analysis showed that fine partitions of the data with >30-50 clusters have limited replicability (Fig. 2h).

To facilitate comparison of additional datasets with ours, we provide helper files, software, and a walkthrough to recapitulate the central results reporting replicability using MetaNeighbor (Supplementary Note). In addition, we demonstrate how the same process of estimating replicability within the BICCN data can be used for cross-comparison and evaluation of novel data.

### Epigenetic cell types of mouse MOp

RNA-Seq data report the cell’s transcriptional state, but do not directly assess the epigenetic modifications of DNA and chromatin configuration that establish and maintain cell identity. Regions of open chromatin and patterns of DNA methylation, including CG and non-CG methylation, are cell type-specific signatures of neuronal identity and can be assayed in single nuclei^16,18^. We applied single-nucleus methylC-Seq (snmC-Seq2^20,35^) and open chromatin (snATAC-Seq^36^) assays to nuclei isolated from the same MOp samples. Independent analyses of each epigenomic dataset identified n=42 cell types from 9,794 cells using snmC-Seq2, and n=33 types from 81,196 cells using snATAC-Seq (Fig. 3; Supplementary Table 4). Marker genes for major cell classes had corresponding patterns of cell type-specific depletion of non-CG methylation (low mCH, Fig. 3b) and open chromatin in the gene body (Fig. 3d).

**Figure 3:**
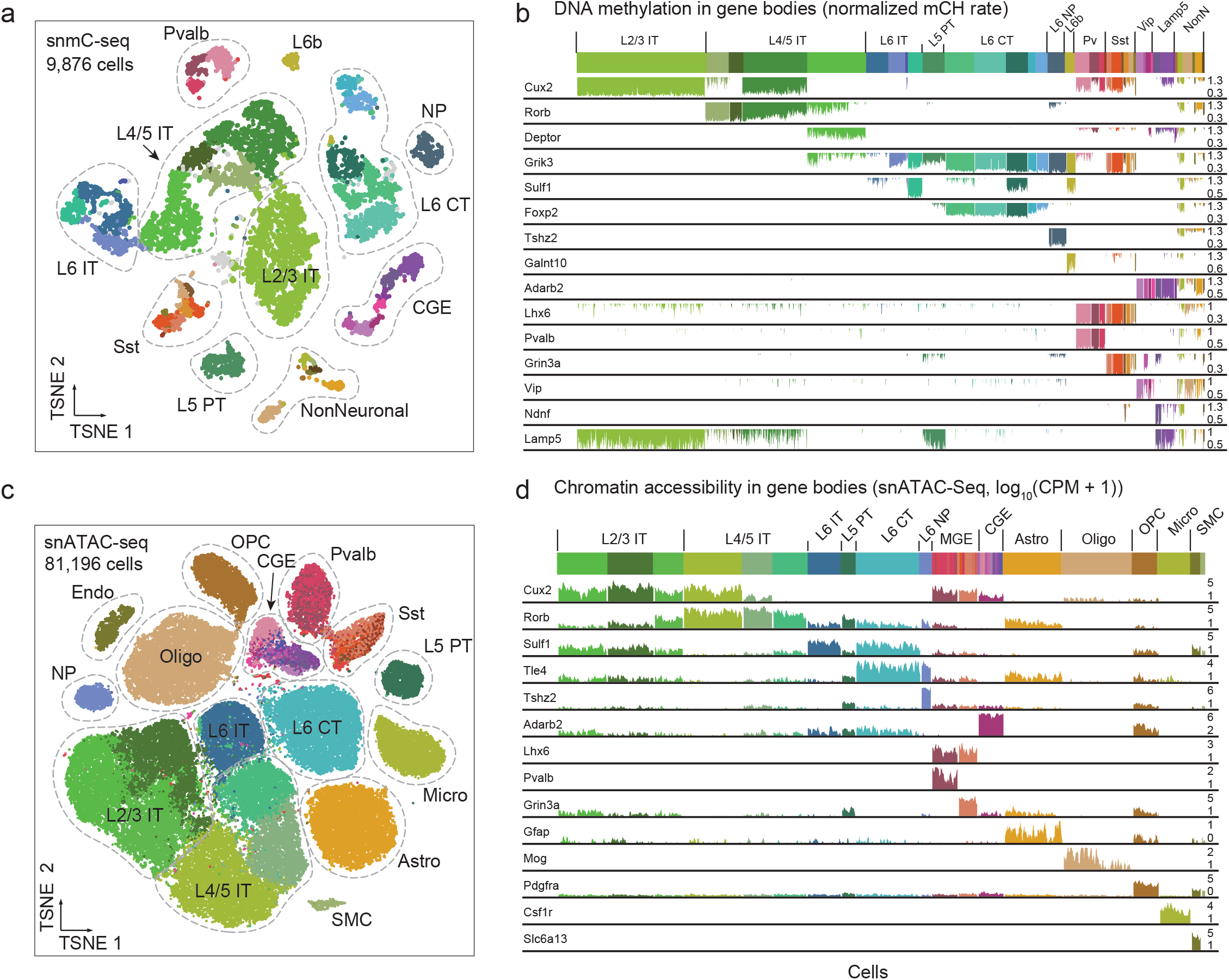
Epigenomic cell types in MOp. **a,** Cell type clusters from single-nucleus methyl-C-Seq (snmC-Seq2^16,20^) for 9,876 MOp nuclei are represented in a two-dimensional projection. Labels indicate broad cell types, colors show finest cluster resolution. **b,** Non-CG DNA methylation level (normalized mCH) for each cell at gene bodies of markers of major cell types. Actively expressed genes have low mCH, indicated by colored bars extending downward. Highly methylated (repressed) genes appear white in this plot. **c,** Two-dimensional projection of cell type clusters from single-nucleus ATAC-Seq (snATAC-Seq^18^) profiles for 81,196 cells. **d,** Gene body chromatin accessibility (total snATAC-Seq read density, log(CPM+1)) for marker genes. For c and d, each bar represents one cell. Cell type abbreviations as in Fig. 2. CGE/MGE - Caudal/Medial ganglionic eminence derived inhibitory cells;

The cell type classifications based on epigenomic datasets were similar to each other and to the transcriptomic classification, despite the significant differences in the biological features assayed, number of cells, genomic coverage, and other parameters. In particular, DNA methylation data provides the highest level of genomic coverage per cell (2.7 million mapped reads on average), similar to the SMART-seq single cell transcriptome datasets (2.1M reads/cell on average). This deep coverage affords precise characterization of cell types using a modest number of cells. To maximize the coverage of DNA methylation in neurons, we applied fluorescence activated nuclei sorting (FANS) to enrich NeuN-expressing cells (95% of collected cells). By contrast, snATAC-Seq generates 8,800 reads per cell but can be applied at a larger scale. For this dataset, no FANS was applied and both neurons and non-neuronal cells were collected (Fig. 1d, Supplementary Table 1).

### Integration of transcriptome and epigenome datasets defines multimodal reference cell types

Although multimodal assays in single cells have been developed^37–39^, the most robust technologies applicable to thousands to hundreds-of-thousands of cells currently rely on destructive measurements that preclude multimodal characterization. Our large-scale census of mouse MOp comprises separate measurements from multiple modalities and technologies. We therefore used computational methods for data integration^37,40–42^ to map the datasets into a common space and to produce a unified, multimodal cell census (Fig. 4, Extended Data Fig. 6). Our overall premise for data integration was that cells of the same type measured in each modality can be identified based on correlated gene-centric features. For example, gene expression is negatively correlated with gene body non-CG methylation^16^ and positively related to the gene body ATAC-Seq read density^38^. Although each dataset differs in systematic ways from the others, these cross-modal correlations allowed us to link cells in each dataset with their most similar counterparts in the other datasets. The eight matched datasets included here, with unprecedented depth and breadth in terms of modalities and technologies, represent a unique opportunity to test the limits of multimodal computational data integration.

**Figure 4:**
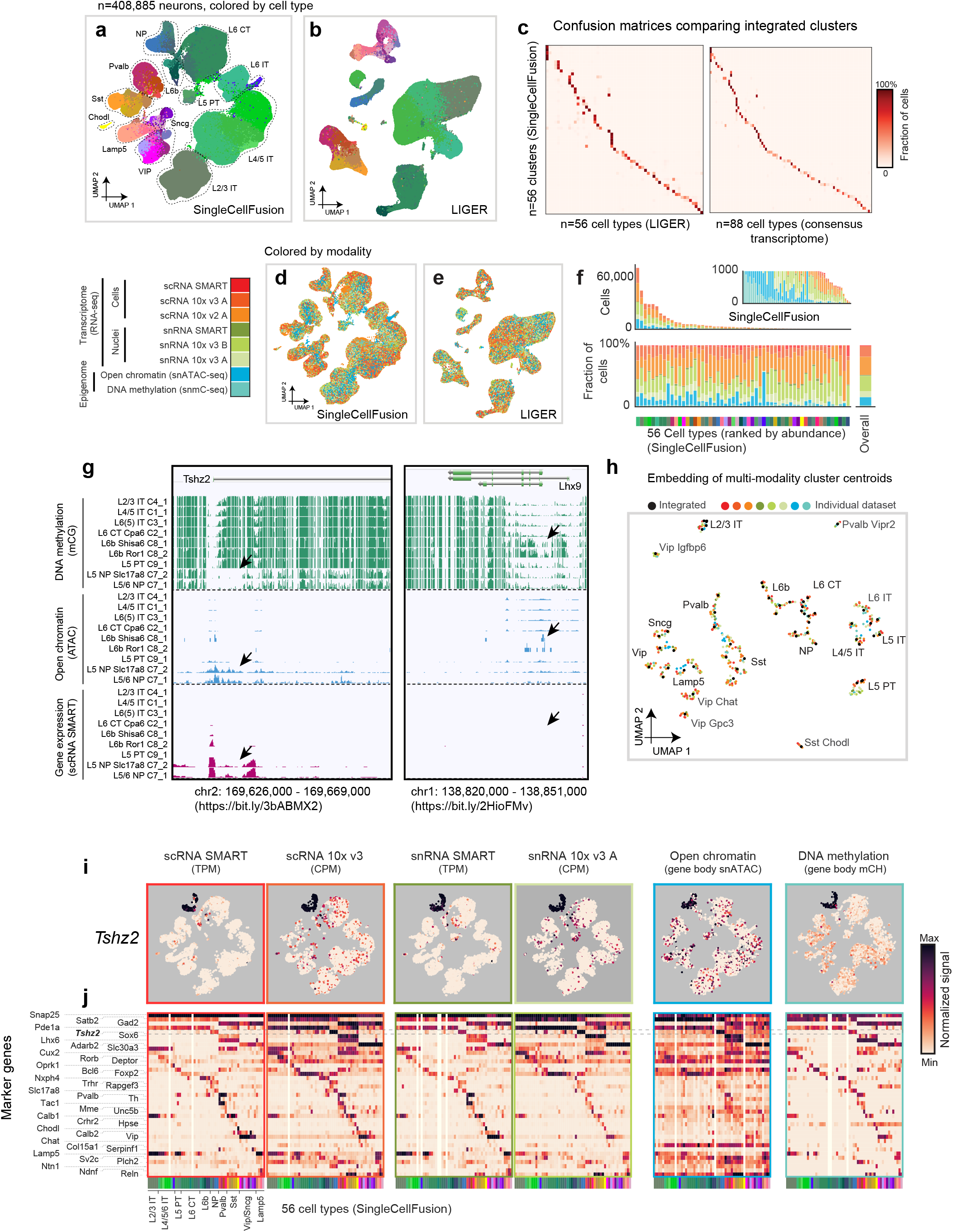
Multimodality integration of >400,000 cells and nuclei. **a,b,** Two-dimensional projection (UMAP) of >400,000 individual cells and nuclei from 8 transcriptomic and epigenomic datasets (excluding snRNA 10x v2 A), integrated using Single Cell Fusion (a) or LIGER (b). Cells are colored by joint clustering assignments from the respective integration method. **c,** Confusion matrices comparing integrated clusters generated by SingleCellFusion versus by LIGER, and comparing SingleCellFusion versus consensus transcriptomic taxonomy. **d,e,** Two-dimensional projection (UMAP) of >400,000 individual cells and nuclei from 8 transcriptomic and epigenomic datasets, integrated using Single Cell Fusion (d) or LIGER (e). Cells are colored by the data modality. **f,** Number of cells in each of 56 multimodality cell types (SingleCellFusion; L2), ranked by cluster size. **g,** Genome browser views across cell types and data modalities. *Tshz2* consistently marks L5 NP cell types across data modalities, whereas Lhx9 marks L6b cell types in DNA methylation signals only. **h,** Embedding of multimodality cluster centroids. Black dots are cluster centroids of integrated clusters (SingleCellFusion); Colored dots are cluster centroids of individual datasets. Cluster centroids are generated by SingleCellFusion. **i,** UMAP embeddings colored by different molecular signals of *Tshz2*. **j,** Heatmaps of marker genes by cell types across data modalities.

The starting point for integration was a set of cell-by-gene matrices summarizing the gene expression, gene-body chromatin accessibility, or gene body non-CG methylation (mCH) for each cell (Supplementary Table 5). We chose to use gene body features to allow us to directly link cells across all three modalities. This strategy does not utilize the cell type-specific epigenetic information outside of gene bodies, including promoters and distal regulatory elements, potentially sacrificing resolution. However, linking distal elements with their associated gene(s) is challenging, and regulatory regions are smaller and thus more affected by sparse coverage in single cell datasets than gene bodies. Although not used for dataset integration, distal regions were subsequently included in the analysis of cell type specific regulation (see below).

We applied two computational approaches based on non-negative matrix factorization (LIGER) and nearest-neighbor imputation (SingleCellFusion)^37,42^ (see Methods; Fig. 4a,b). Both methods identified 56 neuronal cell types, which showed a high degree of concordance between the methods and with the transcriptome-based consensus clusters (Fig. 4c; Extended Data Fig. 6c-f). Gene body-based integration successfully fused all data modalities while preserving fine cell type distinctions (Fig. 4d-f). Indeed, integrated analysis identified more cell types than the single-modality analysis of each epigenomic dataset, while largely concurring with the independent clusters (Extended Data Fig. 6a,b). The data integration was repeated iteratively on 5 major cell classes to provide more interpretable multimodal embeddings (Extended Data Fig. 6c).

After assigning cells to types based on integrated analysis of gene body signatures, we created genome-wide epigenomic and transcriptomic maps for each cell type. By combining the sequencing reads from all cells of a given type for each modality, we generated high-coverage pseudo-bulk data tracks that can be directly compared and analyzed (Fig. 4g). We generated pseudo-bulk tracks at both a relatively coarse (29 cell types, SingleCellFusion level L1) and a fine resolution (56 cell types, SingleCellFusion L2) (Extended Data Fig. 6d). These data can be viewed interactively at https://brainome.ucsd.edu/BICCN_MOp/. Two-dimensional embedding of the centroids of each multimodal cluster together with clusters defined by separate analysis of each dataset shows the close correspondence between the molecular taxonomies (Fig. 4h).

The pseudo-bulk profiles revealed striking examples of cross-modal cell type specific signatures. For example, the *Tshz2* locus is a specific marker of layer 5 near-projecting (NP) excitatory neurons (Fig. 4g), which had low DNA methylation (mCG and mCH), open chromatin, and high levels of cell type-specific gene expression. This gene was identified as a target of the transcription factor *Fezf2* that labels neurons during late-embryonic and early postnatal development^43^. The close correspondence between transcriptomic and epigenomic signatures at *Tshz2,* and 35 markers of other cell types, was evident across each of the datasets (Fig. 4i,j). Importantly, these pseudo-bulk tracks include data, such as CG methylation and intergenic snATAC-Seq signals, that were not used for the multimodal computational integration. The evident alignment of these signatures with the other modalities validates the fidelity of our multimodal clusters.

In addition to concordant cross-modal signals, we also found individual loci where different data modalities did not correspond, suggesting partial decoupling between transcriptomic and epigenomic states. For instance, at the *Lhx9* locus, we found a highly specific enrichment of CG and non-CG DNA methylation in L6b excitatory neurons (Fig. 4g, Extended Data Fig. 6k). *Lhx9* was covered by a large DNA methylation valley (DMV) in each of the other cell types. Despite this cell type-specific epigenetic profile, we found no expression of *Lhx9* RNA in any cell type and only modest enrichment of ATAC-Seq reads. This pattern may represent a vestigial epigenetic signature of embryonic development^44^, as previously described using bulk samples of purified neural populations^17^. Indeed, *Lhx9* has been implicated in early developmental patterning of the caudal forebrain and may be transcriptionally silenced in the adult, potentially via Polycomb-mediated repression^45^. Other regulators of neural development, such as *Pax6* and *Dlx1/2*, have a similar epigenetic profile with cell type-specific hyper-methylation, often accompanied by cell type-specific RNA expression in the hyper-methylated cell type^17,46^.

### Epigenomic signatures of cell type-specific gene regulation

Epigenomic data identify potential regulatory regions, such as distal enhancers, marked by open chromatin and low DNA methylation (mCG). These modalities have complementary technical characteristics, such as the number of cells assayed (higher for open chromatin) and the genomic coverage per cell (higher for DNA methylation; Fig. 1c). We first defined differentially methylated regions (DMRs) and chromatin accessibility peaks independently, identifying 1.49 million DMRs covering 242 Mbp (9% of the genome) and 317,000 accessible regions (ATAC peaks) covering 170 Mbp. In each cell type, a large fraction of accessible regions (35-69%) overlapped with hypo-methylated DMRs, i.e. regions with lower mCG compared to other cell types (Fig. 5a). By contrast, we found many DMRs that did not correspond to accessibility peaks (Fig. 5b). In some cases, we observed that these DMRs coincided with broad open chromatin regions, such as whole gene bodies, which had no narrow ATAC peaks. Notably, we also identified a significant number of accessible peaks which overlapped hyper-methylated DMRs, i.e., regions with higher mCG compared with other cell types (Fig. 5a,b). These regions could indicate regulatory regions bound by methylation-preferring transcription factors^47^.

**Figure 5:**
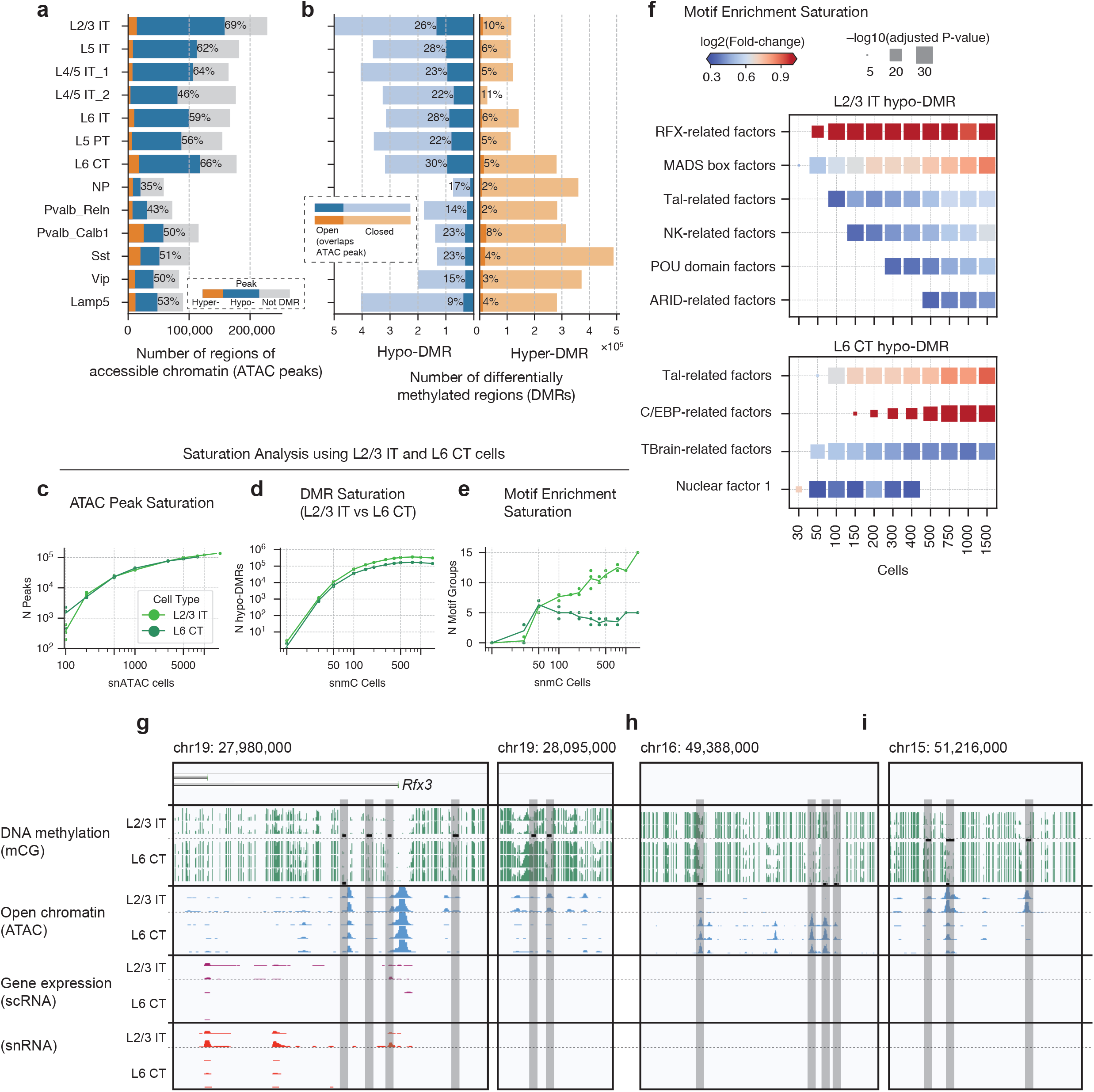
Integrated epigenomic analysis. **a,b,** Thousands of regulatory regions were identified in each cell type using differentially methylated regions (DMRs) and open chromatin regions (ATAC peaks) in multimodal integrated clusters. **c,d** Saturation analysis for two excitatory subclasses shows the number of regulatory regions detected as a function of sampled cells. **e,** Saturation analysis of the number of transcription factor DNA binding sequence (TFBS) motifs enriched in each cell type’s DMRs. **f,** Combining the multimodal information we predicted enhancers using REPTILE^49^, followed by analysis of enriched TFBS motifs. **g-i,** Browser views of loci containing cell type-specific regulatory elements. The *Rfx3* gene is differentially expressed in L2/3 neurons, and has an enhancer specific to L2/3 located -15 kb upstream of the promoter region **(g).** We also found thousands of intergenicregions with accessibility and demethylation specific to L6 CT **(h)** or L2/3 neurons (i).

To assess the comprehensiveness of our regulatory element predictions, we performed saturation analysis taking advantage of the large scale of the integrated data. We focused on two highly abundant subclasses of excitatory neurons, the layer 2/3 intratelencephalic (L2/3 IT, 2 types) and the layer 6 corticothalamic (L6 CT, 3 types) neurons. Each epigenomic dataset includes over 1,000 cells of each of these types. We found that the the number of detectable accessibility peaks increased with the number of sampled cells, without reaching saturation even after sampling 5,000 cells (Fig. 5c). This observation likely reflects the sparse coverage of open chromatin regions in individual cells by snATAC-Seq. By contrast, the number of DMRs for each cell type reached a plateau after sampling 200-300 cells (Fig. 5d).

Cell type-specific enhancers can help to reconstruct regulatory networks, including key transcription factors (TFs) whose binding to DNA at active enhancers may be reflected in the chromatin accessibility and/or DNA methylation signatures. We identified known binding motifs of TF classes^48^ that were enriched in each cell type’s DMRs. Saturation analysis showed that the number of significantly enriched motifs increases with cell number (Fig. 5e), although for L6 CT neurons it reached a plateau of ∼5 key motif families after sampling ∼100 cells. To assess cell type TF networks more comprehensively, we leveraged our integrated DNA methylation (snmC-Seq) and snATAC-Seq data to predict the locations of over 250,000 putative enhancers with fine resolution using machine learning (REPTILE; Supplementary Table 7)^49^. We identified 73,030 putative enhancers in L2/3 neurons, and 66,119 in L6 CT cells. Putative enhancers were distal regions (at least 2 kb from the nearest transcription start site) and, taken together, they represent signatures of the regulatory genome that were not assayed by RNA-Seq (Fig. 5h,i).

Enhancers were enriched in motifs for several TF families^48^ (Fig. 5f). For example, *Rfx* motis were strongly enriched in L2/3 neurons, as previously observed using ATAC-Seq in mouse visual cortex^50^. Using the transcriptomic data, we found that *Rfx3* (but not other *Rfx* family members) was specifically enriched in L2/3 neurons and had substantial gene body hypo-methylation and chromatin accessibility (Fig. 5g). Moreover, we found multiple intergenic regulatory regions with specific signals of open chromatin and low mCG in L2/3 neurons located ∼15 kb upstream of the *Rfx3* promoter. Together with the enrichment of the *Rfx* family binding motif in L2/3 enhancers, these data suggest a key role for *Rfx3* in these neurons. Our findings align with reports of *Rfx3* localization in the superficial portion of L2/3 in mouse somatosensory cortex^51^ and visual cortex^5^.

### Computational validation of cell type reproducibility across datasets

Using our data we sought to define the number of cell types in MOp. Different molecular modalities, sampling strategies and sequencing technologies, as well as different computational analysis procedures, can lead to divergent estimates of the total number of cell types. The difficulty in defining cell types can lead to subjective debate between “lumpers” and “splitters”, hampering progress toward a scientific consensus^9,52,53^. Yet, addressing this question objectively, based on diverse empirical criteria, is essential since it directly determines the granularity of cell types in the cell atlas. In our analyses of MOp transcriptome and epigenome data, we found that many factors could affect the number of derived cell types, from as few as ∼25 cell types to over 100. We therefore pursued a range of analytic methods to cross-validate and assess the statistical and biological reproducibility of cell types. These analyses constrain the range of plausible numbers of cell types based on current single-cell sequencing data. At the same time, they demonstrate that no single estimation of the number of molecularly defined cell types may be objectively supported by currently available methods.

We first addressed the impact of the number of sampled cells on the resolution of the cell atlas. We expect that datasets comprising larger numbers of cells, combined with targeted sampling methods that enrich particularly rare cell types, will saturate the diversity of MOp neurons. Taking advantage of the more than 600,000 sampled cells, we systematically downsampled each dataset and performed community detection with fixed resolution parameter (Fig. 6a). The results showed a logarithmic increase of the number of detected neuronal cell types (clusters) with increased sampling for each of the datasets, with relatively few additional clusters detected after sampling ∼80,000 cells or nuclei. Notably, the dependence of the number of clusters on the number of sampled cells was similar for all modalities and datasets, showing that the number of sampled cells is a key determinant of cluster resolution.

**Figure 6:**
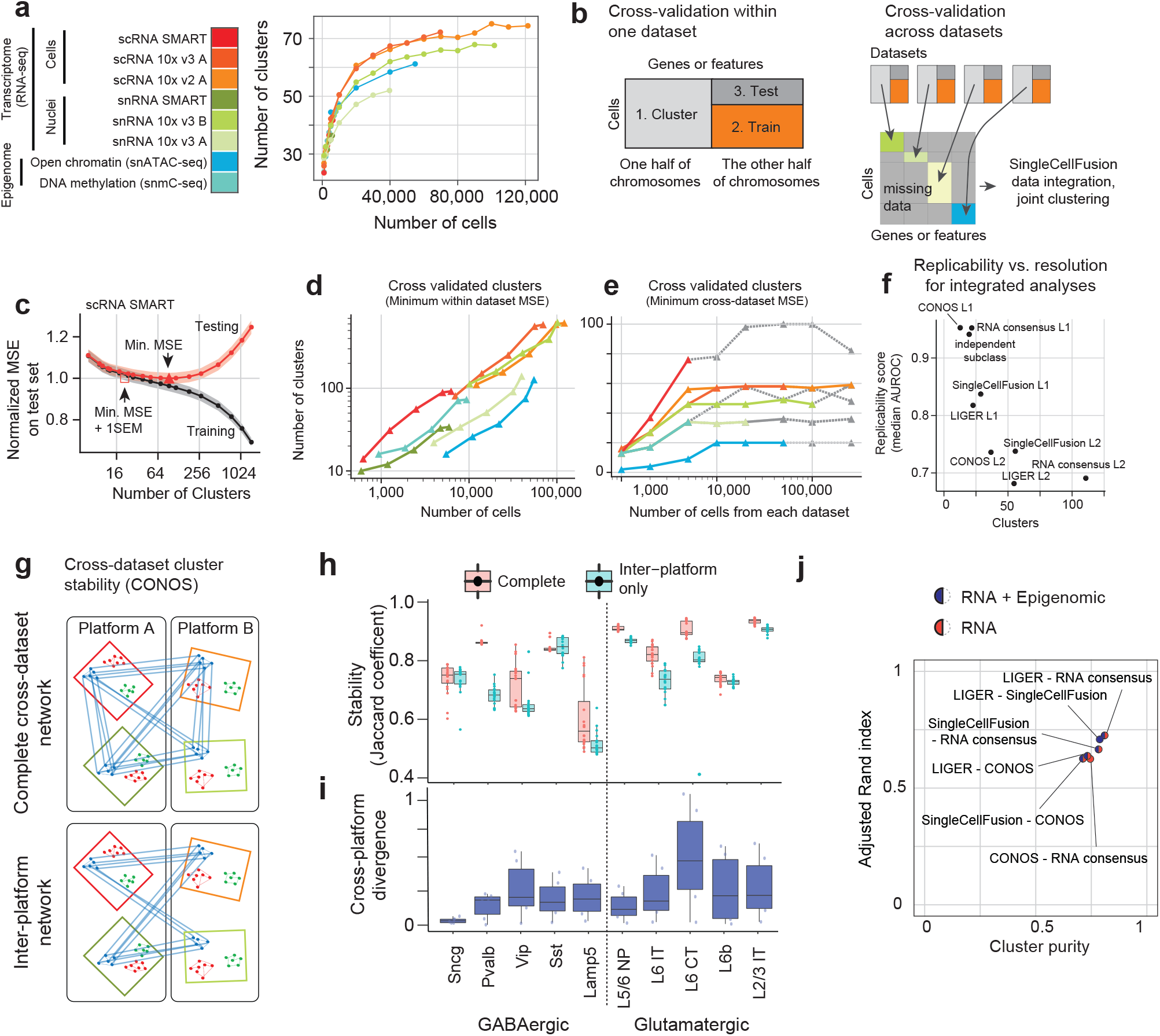
Robustness and reproducibility of cell types within and across datasets. **a,** Number of clusters estimated for each dataset after sampling a fraction of the total cells (Leiden clustering, resolution r=6). **b,** Within- and across-dataset cross-validation scheme. Gene features are split randomly into separate sets for clustering cells (1) and validating the assigned clusters (2,3). After clustering, 80% of cells are used to train a model of the held-out features for each cluster (2). Finally. the remaining cells are used to test model’s prediction on held-out features (3). For cross-dataset comparison, data integration and joint clustering are performed using the first half of genomic features from each dataset, **c,** Test set means squared error (MSE) as a function of the number of clusters obtained by varying cluster resolution for one dataset (scRNA SMART). The minimum MSE and the min. MSE+1SEM defines a range of optimal cluster resolutions outside of which over- and under-clustering lead to poor test-set performance. **d,e,** Number of clusters estimated by within- (d) or across-dataset cross-validation (e), as a function of the number of sampled cells. For cross-dataset comparison, the number of clusters is based on the minimum test MSE for one dataset after joint multimodal clustering, **f,** Trade-off between number of clusters and replicability (median MetaNeighbor AUROC) of consensus clustering methods applied at various resolutions. **g,h** Transcriptomic platform consistency is assessed by cross-dataset cluster stability analysis (CONOS) using complete networks. and using inter-platform edges only. Glutamatergic and Pvalb subclasses have reduced stability in inter-platform comparison, **i,** Cross-platform expression divergence (Jensen-Shannon) for major cell subclasses. **j,** Agreement between consensus clustering results using different computational procedures.

Any dataset can be divided into increasingly fine-grained clusters, but those clusters may not reflect biologically meaningful or reproducible cell type distinctions. We therefore devised cross-validation schemes to objectively measure the generalizability of cluster-based descriptions of the data (Fig. 6b). We first used within-dataset cross-validation, dividing the set of genomic features into two parts (for clustering and validation, respectively). After clustering all cells, we then split the cells into training and test sets. By training a classifier to predict the validation features using the cluster labels, and applying this classifier to the test set cells, we could measure the mean squared error (MSE) of our cluster-based prediction of single cell transcriptomic or epigenomic features in the test set. We applied this procedure to each dataset with a range of clustering resolutions, resulting in a U-shaped cross-validation curve for test set error as a function of the number of clusters (Fig. 6c). The location of the minimum MSE is an estimate of the number of reliable clusters. Finally, we repeated this cross-validation procedure for each dataset in combination with systematic downsampling (Fig. 6d).

This analysis highlights the different depth of information per cell from each modality. Notably, all of the datasets (except snRNA SMART-seq) supported ∼100 or more cell types when a sufficient number of cells were sampled, although the number of cells required was larger for snATAC-Seq compared with RNA-Seq or snmC-Seq. The latter is understandable given the sparseness of the snATAC-seq data. We further found that sc/snRNA-Seq datasets with the largest numbers of cells could support very high cluster resolution with up to ∼600 clusters. Our cross-validation analysis shows that these fine-grained clusters capture genuine transcriptomic structure which is correlated and replicable across cells and across genomic features. However, it is likely that at least some of this structure corresponds to continuous variation within discrete cell types, rather than truly discrete cell type categories^54^. Moreover, the cross-validation analysis shows that there is no sharply defined error minimum at a particular value of the number of clusters. Instead, the U-shaped cross-validation curve has a broad basin covering a range of plausible values (Fig. 6c).

To more stringently test the reproducibility of cell types, we performed cross-dataset cross-validation (Fig. 6b; Methods). This procedure uses a randomly chosen half of genomic features to perform data integration and joint analysis of eight datasets using SingleCellFusion. Next, we use the joint cluster labels to perform cross-validation in each dataset, as in the within-dataset procedure above. This analysis supported a maximum resolution of ∼100 clusters when testing using the scRNA SMART-seq data (Fig. 6e).

As an alternative to joint analysis of multiple datasets, which could potentially discern spurious correlations due to computational data integration, we also took a more stringent approach to cross-validation. Using the independent cluster analysis of each dataset as an input, we performed MetaNeighbor analysis to assess the replicability of clusters^31^. We found that the median replicability score for all clusters was very high (AUROC > 0.8) for integrated analyses with coarse resolution (<50 clusters, level 1 (L1) analyses; Fig. 6f). The more fine-grained joint analyses (L2, 50-120 clusters) were also largely supported by MetaNeighbor, but with a lower median replicability score around 0.7. Notably, we found a high degree of consistency in the results of joint cluster analysis when using different computational methods (Fig. 6j).

Finally, we explored whether MOp cell type signatures were largely stable across different sc/snRNA-Seq platforms. Using four RNA-Seq datasets (scRNA SMART, snRNA SMART, scRNA 10x v3 A, and snRNA 10x v3 A), we performed clustering on network of samples (CONOS^55^) to link cells across datasets and determine joint clusters. We compared the clustering results based on inter-platform network connections only vs. results that also included connections across datasets of the same platform (Fig. 6g). Most neuron types, with the exception of Pvalb and L6 CT, had the same level of cluster stability (as assessed by bootstrap sampling of cells) using both approaches (Fig. 6h) and a low level of inter-platform divergence in their cell type transcriptomic signatures (Fig. 6i).

## Discussion

Our mouse primary motor cortex (MOp) atlas represents the most comprehensive, integrated collection of single cell transcriptome and epigenome datasets for a single brain region to this date. We generated a high resolution consensus transcriptomic cell type taxonomy that integrates seven sc/snRNA-Seq datasets collected from MOp with six experimental methods. Our MOp transcriptomic taxonomy is highly consistent with a previously published transcriptomic cell census from VISp/ALM based on SMART-Seq alone^5^. We found that gene expression profiles are largely consistent across different methodologies, while providing complementary information about particular genes such as nucleus-enriched transcripts. We find molecular signatures of putative L4 excitatory neurons^26^, as well as multiple types of L5 PT neurons that align with recently described populations with distinct subcortical projection targets^24^. The MOp atlas demonstrates the power of a two-pronged strategy that uses broad-sampling of diverse cell types (e.g. 10x with large number of cells and shallow sequencing) together with deep-sequencing (e.g. SMART-Seq) to precisely characterize gene expression profiles for each cell type. These insights should guide future cell census efforts, by the BICCN and others, at the scale of whole brains and in other species.

Going beyond RNA sequencing, we further demonstrate multimodal integration of transcriptome (sc/snRNA-Seq), DNA methylation (snmC-Seq2), and chromatin accessibility (snATAC-Seq) datasets using two computational methods (SingleCellFusion and LIGER). It is possible to directly establish links between molecular modalities through simultaneous measurement of multiple signatures in the same cell^56,57^. However, multimodal single-cell assays remain challenging and often sacrifice the depth or resolution of data in each modality compared with single modality assays. Moreover, it is important to show that data collected from different individual animals, across different laboratories and using different experimental platforms and assays, nevertheless can be integrated within a unified cell type atlas. By correlating mRNA transcripts, gene-body methylation and accessibility peaks, we showed that different types of data can be integrated without sacrificing the resolution of >50 fine-grained neuron types. Our integrated data link cell type-specific transcription with hundreds of thousands of cell type-specific regulatory elements including distal enhancers. Combining transcriptional and epigenetic signatures of cell identity will enable development of new tools for cell targeting and manipulation utilizing newly discovered cell type-specific promoters and enhancers.

We took advantage of the unprecedented diversity of large-scale datasets, collected in coordinated fashion from mouse MOp, to critically evaluate the robustness and reliability of the cell type taxonomies obtained by clustering of various molecular datasets. Using MetaNeighbor^31^ and CONOS^55^, we quantified the reproducibility of cell types across independent RNA-Seq datasets and analyses and found 70 clusters with high reproducibility. These data demonstrate the tradeoff between highly reproducible, coarse-grained classifications at the level of cell classes, and fine-grained classifications of cell types which may be less statistically and/or biologically reproducible. Our cross-validation analysis of individual datasets and multimodal integration objectively constrains the range of cluster resolutions supported by the data without overfitting. Rather than supporting a single, definitive number of cell types in mouse MOp, our studies instead converge on the conclusion that a range of cluster resolutions spanning from ∼30 to as many as 116 cell types is supported by the data. Indeed, discrete cell type categories may be an inappropriate description at a fine-grained level of analysis, where the cells’ molecular profiles vary along a continuum. Cross-modality integration and analysis of cluster reproducibility can constrain the appropriate range of cluster resolutions, and can also reveal the features of the cell type taxonomy that are supported across multiple biological features. Progress in understanding the functional transcriptional signatures that shape cell identity and granularity may further clarify cell type classification^53^. Overall, the data and analyses presented here support the classification of at least 55 neuron types in the mouse MOp, forming a complex landscape of cellular diversity.

By integrating nine large-scale single cell transcriptome and epigenome datasets, we have comprehensively classified and annotated the diversity of cell types in the adult mouse primary motor cortex (MOp). Our study demonstrates general procedures for objective cross-dataset comparison and statistical reproducibility analysis, as well as standards and best practices that can be adopted for future large-scale studies. Together with complementary BICCN datasets from spatial transcriptomics, connectivity and physiology, as well as cross-species comparative studies, our results help to establish a multi-faceted understanding of brain cell diversity. Targeted studies of individual cell types, taking advantage of the transcriptional and epigenetic signatures described here, will define their functional roles and significance in the context of neural circuits and behavior. Integrative analyses will be essential to make progress toward an encyclopedic atlas of brain cell types that distills the essential organizational structure reflected in diverse molecular signatures.

## Acknowledgments

We are grateful to Anita Bandrowski and Yong Yao for insightful comments. This work was funded by the NIH BRAIN Initiative (U19MH114830 to H.Z.; U19MH121282 to J.R.E.; U19MH114821 to Z.J.H.; R24MH114788 to O.R.W.; U24MH114827 to M.H.; R24MH114815 to R.H./O.R.W.; NIH NIDCD DC013817 to R.H.), the Hearing Restoration project Hearing Health Foundation (R.H.), and NIH NIGMS (GM114267 to H.C.B.).

## Author contributions

Contribution to RNA data generation: A.R., A.T., B.T., C.R., C.R.V., D.B., D.M., E.L.D., E.Z.M., H.T., H.Z., J.G., J.S., K.C., K.L., K.S., M.K., M.T., N.D., N.M.N., O.F., T.C., T.N.N., T.P.

Contribution to mC data generation: A.B., A.C.R., A.I.A., A.P., C.L., H.L., J.D.L., J.K.O., J.R.E., J.R.N., M.M.B., S.N., Y.E.L.

Contribution to ATAC data generation: A.P., B.R., J.D.L., J.K.O., M.M.B., S.P., X.H., X.W., Y.E.L.

Contribution to data archive/infrastructure: A.M., B.R.H., C.C., C.V.V., E.A.M., F.X., H.C., H.C.B., J.C., J.G., J.K., J.O., M.G., M.H., O.R.W., R.F., R.H., R.S.A., S.A.A., S.N., V.F., W.I.D., Z.Y.

Contribution to data analysis: A.R., A.S.B., B.T., D.R., E.A.M., E.D.V., E.P., E.Z.M., F.X., H.L., H.R.D.B., H.Z., J.D.W., J.G., J.G., J.O., K.S., K.S., K.V.D.B., L.P., M.C., O.F., O.P., P.V.K., Q.H., R.F., S.D., S.F., S.N., T.B., V.N., V.S., W.I.D., Y.E.L., Z.Y.

Contribution to data interpretation: A.R., B.R., B.T., C.L., E.A.M., E.D.V., E.Z.M., F.X., H.L., H.Z., J.D.W., J.G., J.N., M.C., M.M.B., P.V.K., Q.H., R.F., S.F., T.B., Y.E.L., Z.Y.

Contribution to writing manuscript: A.S.B., E.A.M., F.X., H.L., H.Z., J.D.W., J.G., L.P., M.C., Q.H., S.F., Z.J.H., Z.Y.

## Data access and analysis resource

The BICCN MOp data (RRID:SCR_015820) can be accessed via the NeMO archive (RRID:SCR_002001) at accession: https://assets.nemoarchive.org/dat-ch1nqb7. Visualization and analysis resources: NeMO analytics: https://nemoanalytics.org/, Genome browser: https://brainome.ucsd.edu/annoj/BICCN_MOp/, Epiviz browser: https://epiviz.nemoanalytics.org/biccn_mop

---

**Extended Data Figure 1:**
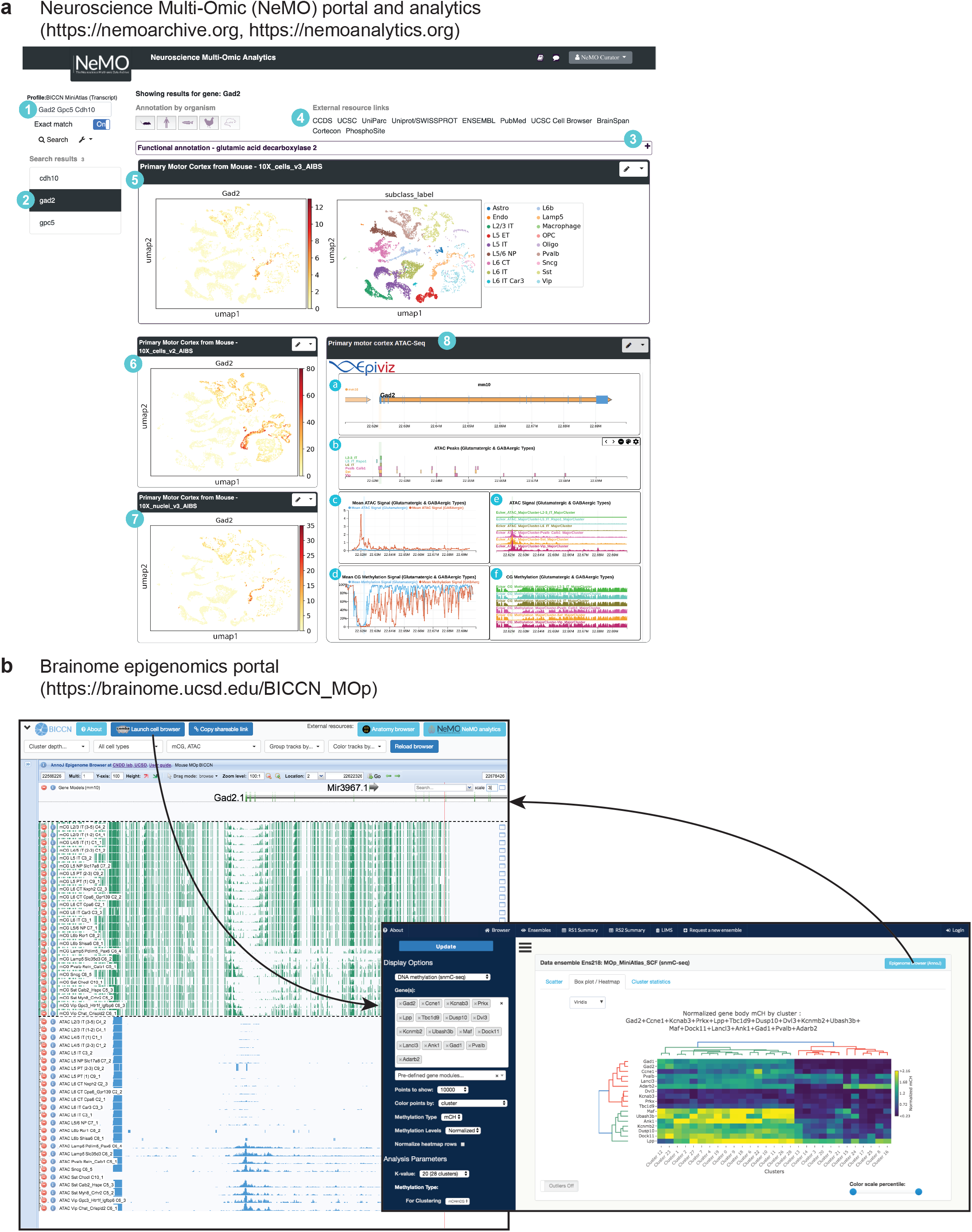
Interactive data access, visualization, and analysis. **a,** NeMO Analytics (nemoanalytics.org) visualization and analysis environment for the BICCN mouse molecular mini-atlas. Screenshot of NeMO Analytics showing multi-omic results for glutamate decarboxylase 2 (*Gad2*), a marker gene in inhibitory neurons. The web portal has the following features: (1) Search box for gene names; (2) Indicator of gene viewed; (3) Expandable species-specific functional annotation; (4) Link-outs to additional resources for the selected gene; (5,6,7) interactive visualizations of each BICCN dataset, displayed in a ‘standalone’ box showing gene expression and cell clustering on integrated UMAP coordinates. Additional data exploration options for each of the datasets are available via the drop-down menu at the upper right corner of the NeMO Analytics dataset titles. (8) An embedded Epiviz interactive workspace to visualize scATAC-seq and sncMethyl-seq datasets in a linear browser view (a), here showing the average ATAC and % CG methylation at the *Gad2* locus (c,d) as well as in each major cluster of glutamatergic and GABAergic neurons (b,e,f). Epigenomic data are also available at http://epiviz.nemoanalytics.org/biccn_mop, and instructions for setting up and extending the Epiviz workspaces are available at http://github.com/epiviz/miniatlas. **b,** Brainome epigenomics portal (brainome.ucsd.edu/BICCN_MOp). The portal shows single base resolution epigenomic and transcriptomic data (snmC-Seq, snATAC-Seq, sc/snRNA-Seq) using the AnnoJ browser. Drop-down menus allow the user to select groups of cells (e.g. Excitatory, Inhibitory, MGE-Derived, etc.), modalities (mCG, mCA, ATAC, scRNA, snRNA, enhancers), and display options. A Cell Browser allows visualizing scatter plots and heatmaps of groups of genes across data modalities.

**Extended Data Figure 2:**
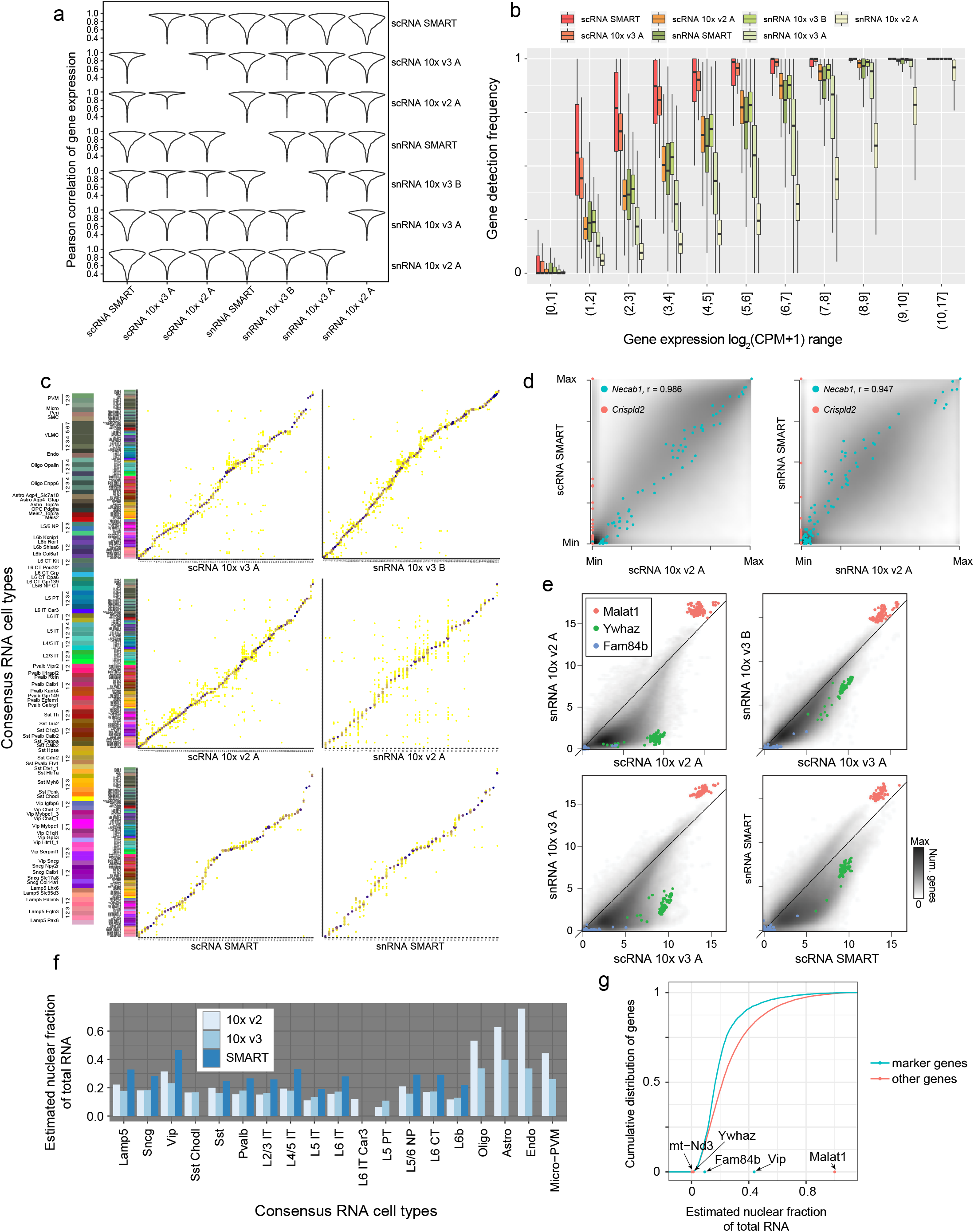
Cluster membership and gene expression consistency across sc/snRNA-Seq datasets. **a,** Pearson correlation of gene expression of 3,792 cell type-specific marker genes across cell types between every pair of datasets. Each violin plot shows the distribution of correlation values for all genes between a pair of datasets. Most genes have highly conserved gene expression patterns at cell type level among all datasets (average correlation 0.856 across all pairs of comparisons). The most consistent datasets are scRNA 10x v2 and v3 (average correlation 0.95), while snRNA 10x v3 B is also highly similar to both scRNA 10x v2 and v3 datasets. Overall, we found the differences between single cell and single nucleus datasets to be more significant than SMART-seq versus 10x platform differences. **b,** Gene detection frequency (sensitivity) at each gene expression range for each dataset. Expression of all genes in each cell type was binned based on the average logCPM in scRNA 10x v2 and snRNA 10x v3 B datasets. Single cell datasets overall have higher sensitivity for gene expression than single nucleus datasets, with the exception of snRNA 10x v3 B dataset, which was more sensitive than scRNA 10x v2 A dataset. For weakly expressed genes, the gene detection frequency can vary dramatically between datasets. For these genes, scRNA SMART was the most sensitive, followed by 10x v3 datasets, all of which showed very robust gene detection. Note that sequencing depth was not considered for this analysis. **c,** Comparisons between clustering analysis of individual datasets with the consensus clusters derived from seven transcriptome datasets. The size of the dot indicates the number of overlapping cells, and the color of the dot indicates the Jaccard index (number of cells in intersection/number of cells in union) between the independent and joint clusters. **d,** Comparison of the relative gene expression of marker genes across all cell types between corresponding SMART-seq and 10x v2 datasets. To compare gene expression directly between SMART-seq and 10x datasets, which differ in experimental platforms, gene expression quantification software and gene annotation reference, for each gene, we normalized the average log2(CPM+1) values at the cluster level in the range [0,1] by subtracting the minimum value and then dividing by the maximum value for that gene. The smooth scatter plot corresponds to the normalized gene expression for all marker genes across all types in two datasets, with their overall Pearson correlation (across all marker genes and cell types) highlighted. **e,** Differential enrichment of transcripts in single cells (x-axis) vs. single nuclei (y-axis) across four platforms. Axis labels are the same as in Fig. 2f. Non-coding RNAs such as *Malat1* are enriched in nuclei. **f,** Using the ratio of *Malat1* expression between corresponding sn/scRNA-seq datasets, we estimated the fraction of nuclear content for each subclass as described previously^28^. snRNA 10x v3 B dataset is not used for this estimate as it also captures cytoplasmic mRNAs according to e. If a dataset includes <20 cells or nuclei in a given subclass, then the corresponding pair is not shown. For cells with large somata, such as L5 PT cells, snRNA datasets only capture 5-20% of the mRNAs of the corresponding scRNA datasets. For cells with smaller somata, such as glia, the ratio is larger, suggesting most of the mRNAs are nuclear. **g,** Distribution of the estimated nuclear localization fraction for all mRNAs based on comparison of the sn/scRNA 10x v2 datasets^28^. To calibrate the differences among cell types, we sampled the same number of cells in each cluster for both datasets, and aggregated all the cells for estimation. We plot the empirical cumulative density function for the marker genes and all other genes separately. The fraction of nuclear mRNAs for five selected genes are shown along the X axis. As expected, mitochondrial genes such as *mt-Nd3* have almost no nuclear localization, while *Vip* is significantly enriched in the nucleus. A selected set of 3,792 cell type-specific marker genes (see Methods section “Marker gene selection”) have lower nuclear fraction relative to the other genes (median 16.6%, compared with 21.9% for non-marker genes).

**Extended Data Figure 3:**
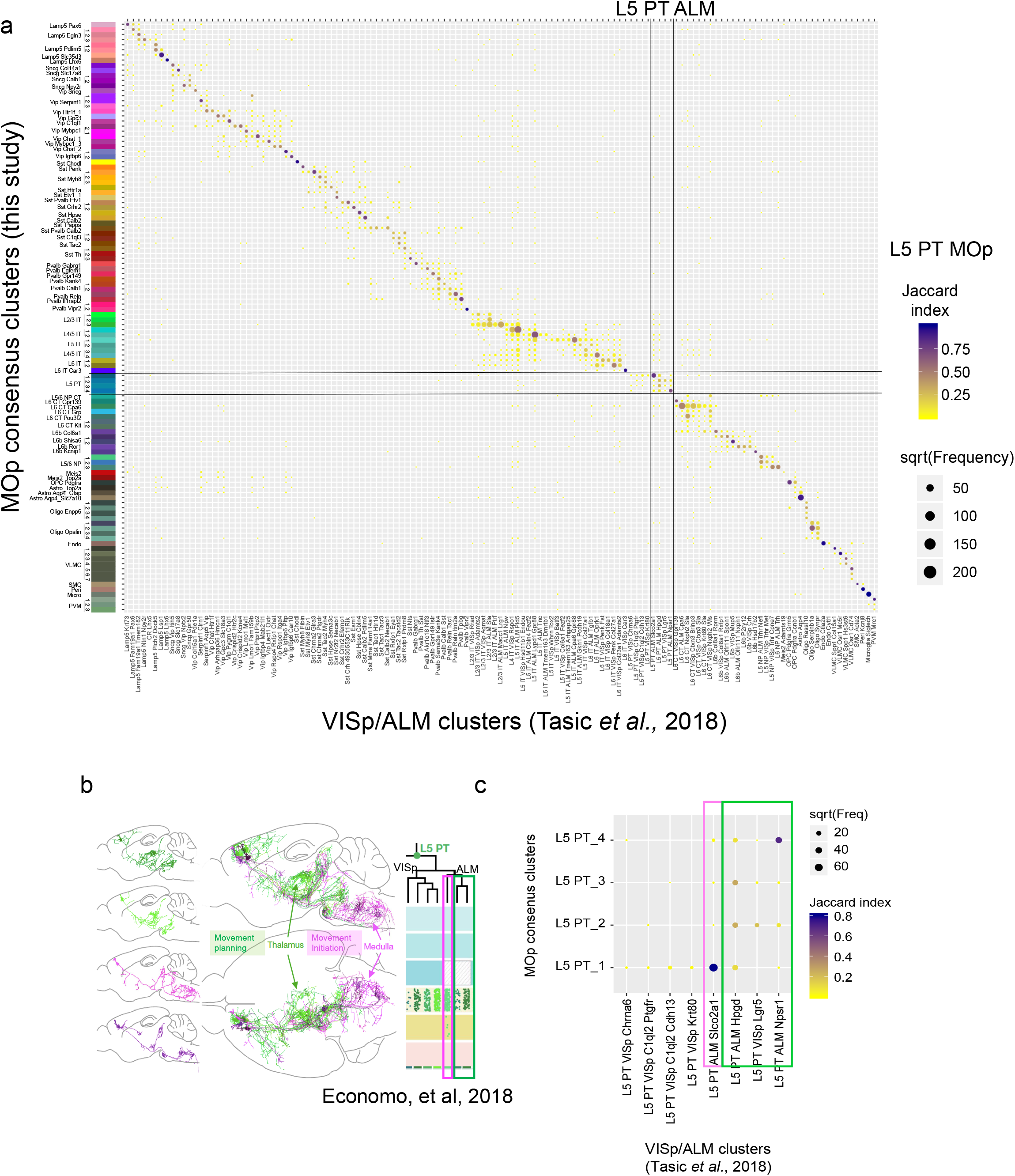
Correspondence between MOp consensus RNA-Seq cell type taxonomy and previously published VISp/ALM cell type taxonomy^5^. **a,** Cells from all sc/snRNA MOp datasets were mapped to the most correlated VISp/ALM cell types based on VISp/ALM cell type markers. The size of dots indicates the number of overlapping cells, and the color indicates the Jaccard index (number of cells in intersection/number of cells in union). MOp L5 PT types are mapped predominantly to L5 PT ALM types in the VISp/ALM study. **b,** Three L5 PT ALM types can be divided into two groups with distinct projection patterns. Cells in the pink group project to medulla and have been functionally associated with movement initiation, while the cells in the green group project to thalamus, associated with movement planning. Adapted from (Economo, et al. 2018)^24^. **c,** Enlarged view of the correspondence between MOp and VISp/ALM L5 PT types. Two subsets of medulla-projecting (pink) and thalamus-projecting (green) L5 PT cells are highlighted.

**Extended Data Figure 4:**
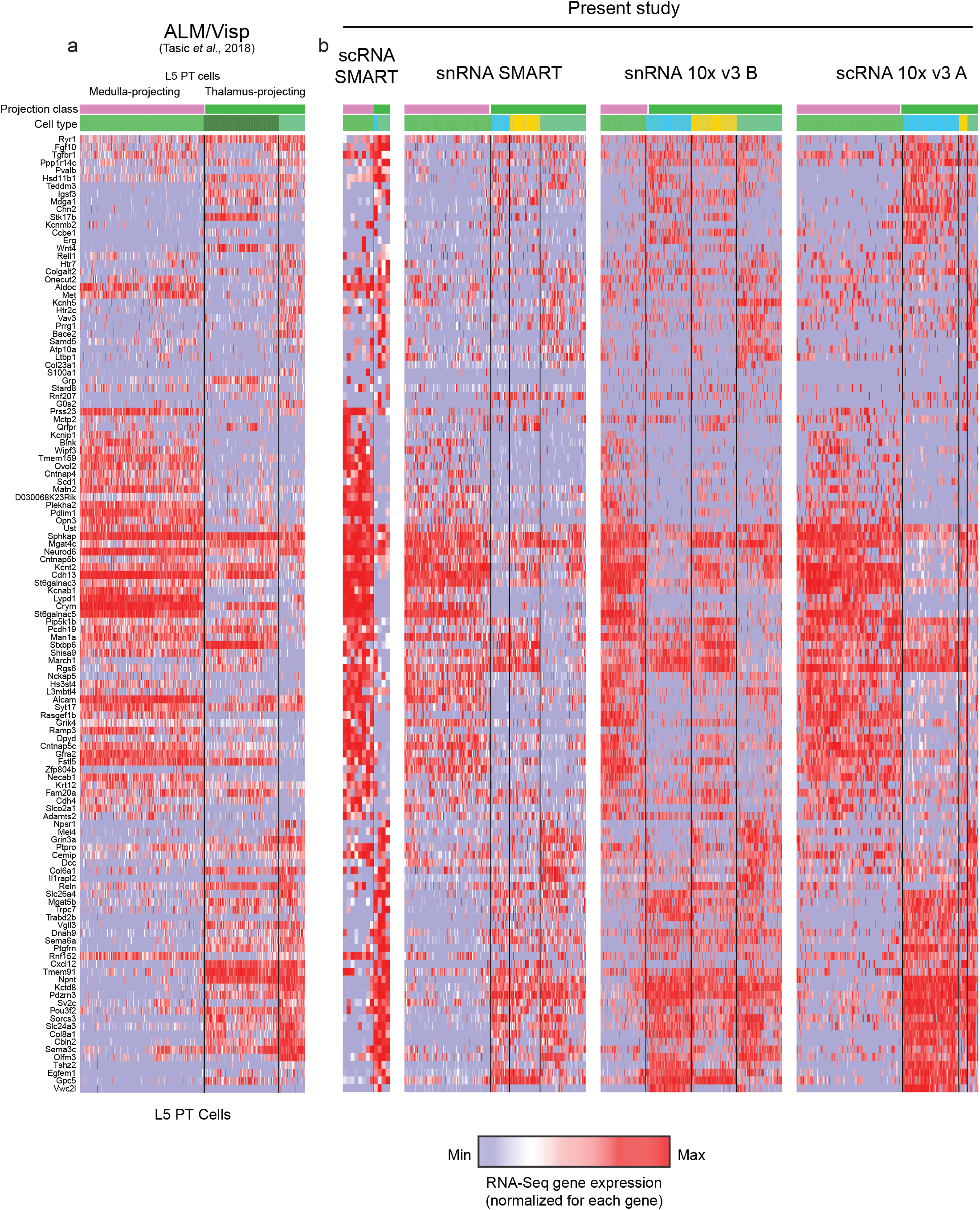
Marker genes for L5 PT cell types. **a** Heatmap showing expression of a combination of marker genes of L5 PT ALM types in previously published dataset^5^, and marker genes for MOp L5 PT types The color bars on the top indicate the cell type and projection class. **b,** Heatmap for MOp L5 PT types in multiple sc/snRNA datasets using the same marker genes in the same order as in a. Cell types are divided into the pink and green groups based on correspondence in Extended Data Fig. 3c.

**Extended Data Figure 5:**
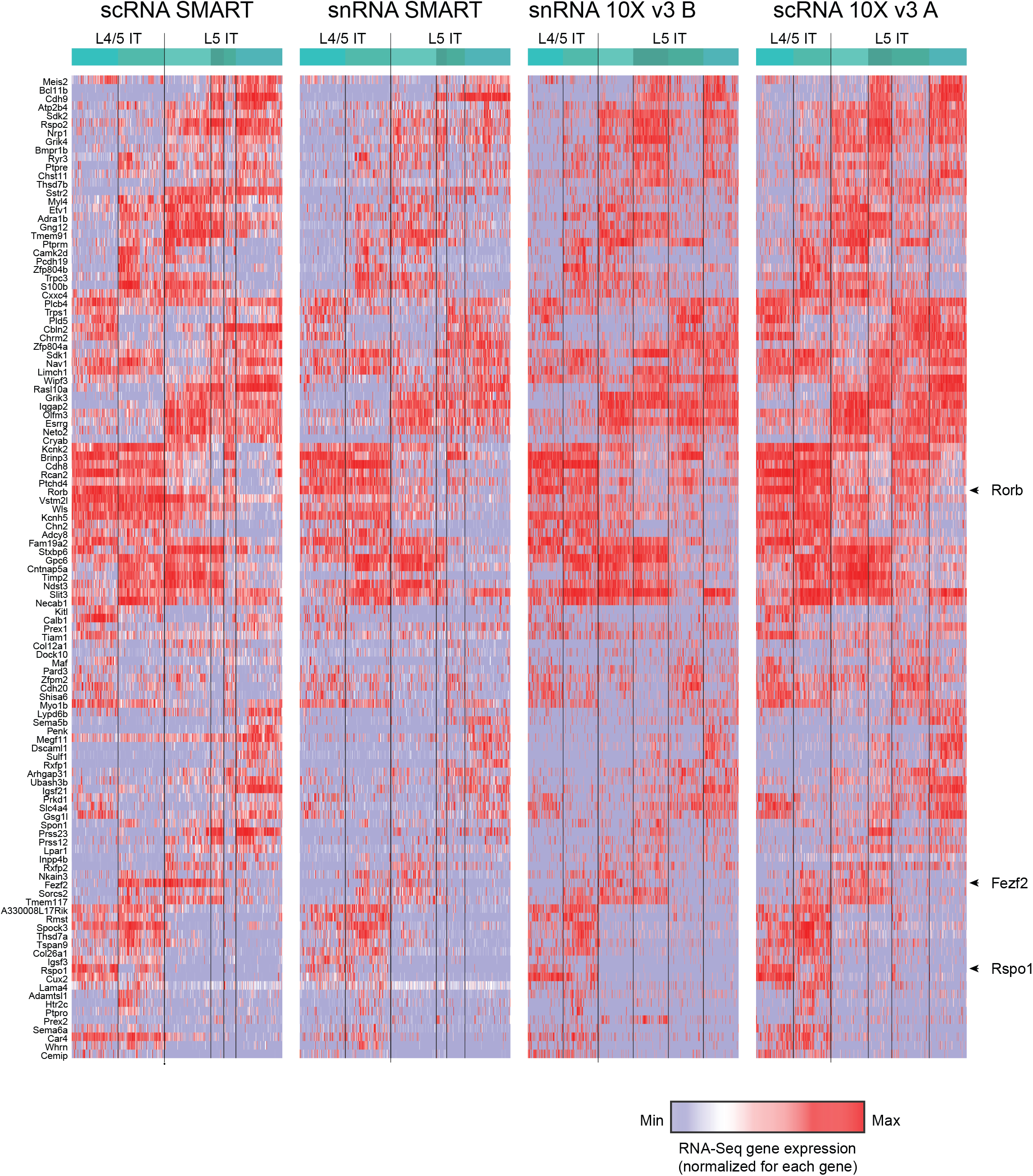
Marker genes for L4/5 IT and L5 IT cell types. **a,** Heatmap showing expression of a combination of marker genes of L5 PT ALM types in previously published dataset^5^, and marker genes for MOp L5 PT types The color bars on the top indicate the cell type and projection class. **b,** Heatmap for MOp L5 PT types in multiple sc/snRNA datasets using the same marker genes in the same order as in a. Cell types are divided into the pink and green groups based on correspondence in Extended Data Fig. 3c.

**Extended Data Figure 6:**
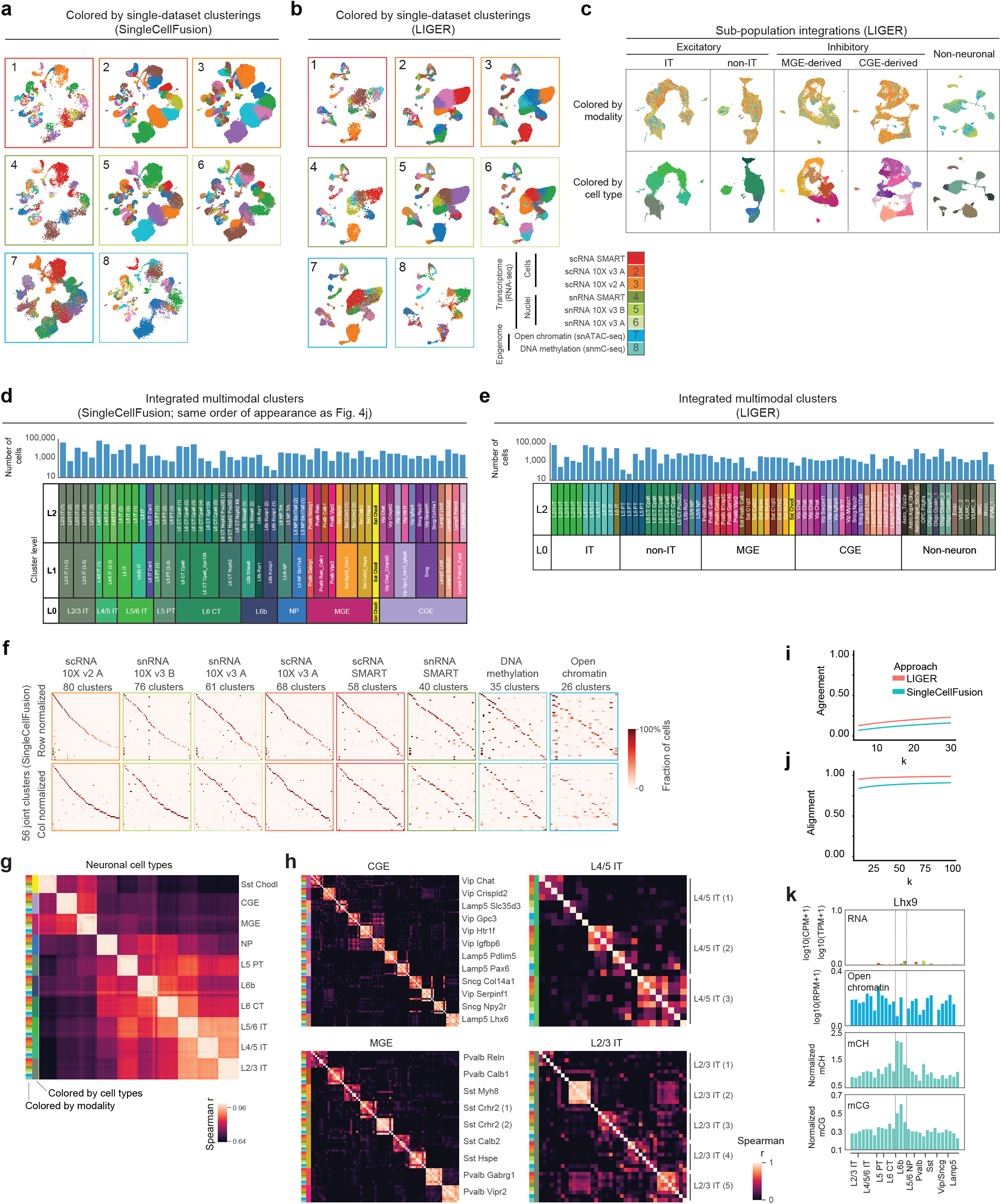
Validation of multimodal integration of transcriptomic and epigenomic data. **a,b,** Integrated, multimodal UMAP embeddings (a: SingleCellFusion; b: LIGER) colored by the clusters assigned in separate analysis of each dataset. Each panel shows the cells from a single dataset. **c,** Integrated analysis of major cell classes by LIGER. Cells in each of 5 cell classes are separately integrated, illustrating fine-grained resolution of integrated data. **d,e** The number of cells for 56 integrated clusters (d: SingleCellFusion L2; e: LIGER L2), as well as the corresponding coarser clusters (L1, L0). Cluster order and color scheme are the same as shown in Fig. 4a,j. **f**, Confusion matrix comparing integrated clusters (SingleCellFusion L2) with single-modality clustering for every dataset. **g**, Spearman correlation matrix for cluster centroid gene expression (measured or imputed) across major cell subclasses for each dataset (SingleCellFusion L0). **h**, Correlation for subsets of inhibitory (CGE, MGE) and excitatory (L4/5 IT, L2/3 IT) neuron types using fine-grained integrated clusters (SingleCellFusion L2). **i,j,** Agreement and alignment metrics^42^ characterize the fidelity of the joint low-dimensional embedding for LIGER and SingleCellFusion. Agreement measures the fraction of k-nearest neighbors for each dataset are still nearest neighbors in the low-dimensional embedding. A high value of the agreement metric thus indicates preservation of each dataset’s internal structure in the joint embedding. Alignment measures the mixing of datasets in the joint low-dimensional space, and is a normalized measure of the mean number of k-nearest neighbors that come from each of the datasets. **k,** Multimodal molecular signals of the developmentally expressed gene *Lhx9* across cell types (n=29; SingleCellFusion L1), showing specific accumulation of mCG and mCH in L6b neurons with no corresponding RNA or ATAC-Seq signal.

**Extended Data Figure 7:**
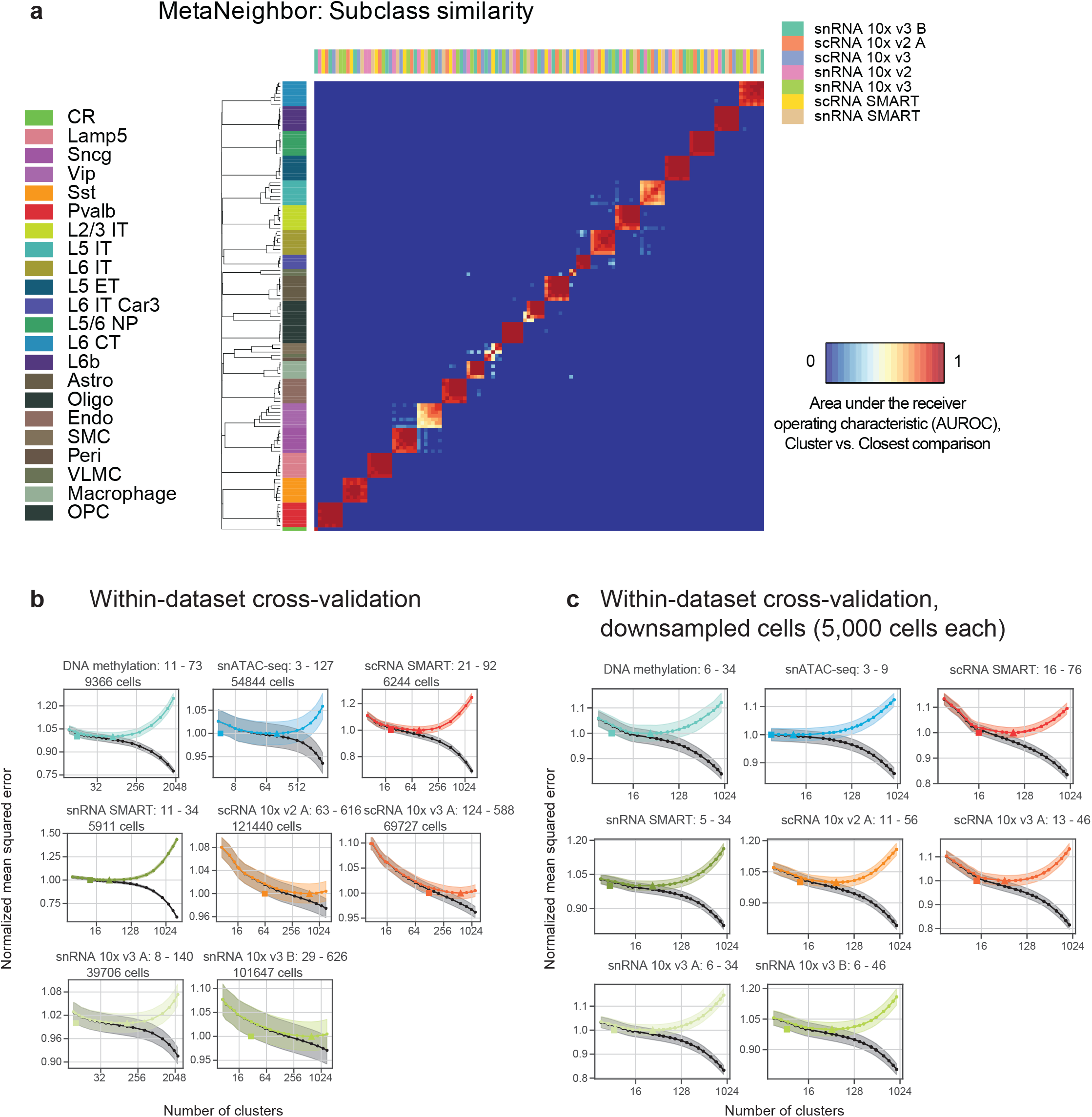
Metaneighbor and cross-validation analysis of cluster reproducibility. **a,** Heatmap showing replicability scores (MetaNeighbor AUROC) at the subclass level of the independent clusterings of seven RNA-Seq datasets. High AUROC indicates that the cell type labels in one dataset can be reliably predicted based on the nearest neighbors of those cells in another dataset, together with the independent cluster analysis of that dataset. **b,c** Within-dataset cross-validation analysis for each dataset, either using the full set of cells (b) or using a random sample of 5000 cells (c). In each plot, the black curve shows training error while the colored U-shaped curve shows the test set error, with a minimum at the cluster resolution that balances over- and under-fitting.

**Extended Data Figure 8:**
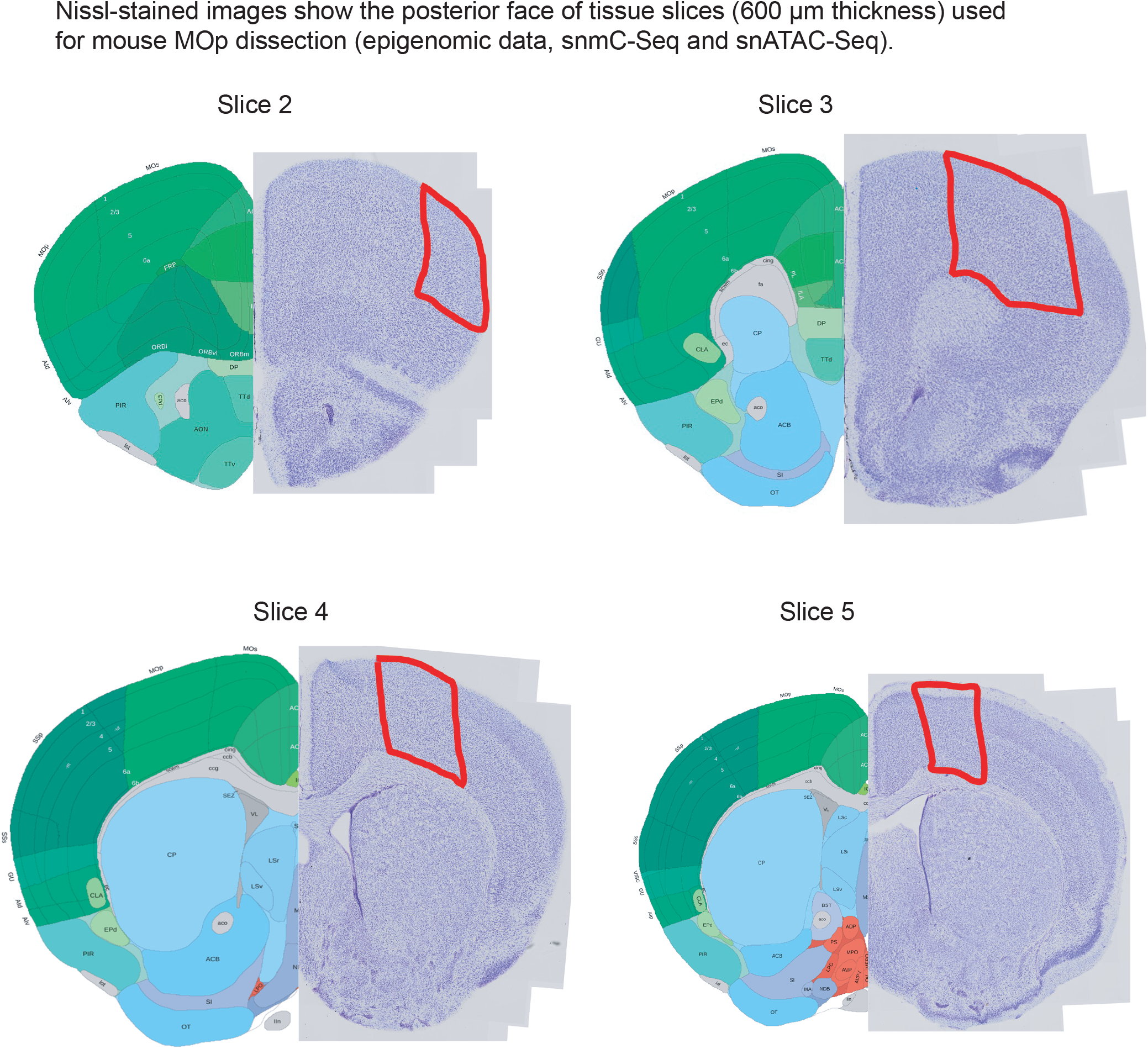
Diagrams of brain slices and dissected regions for epigenomic data samples (snATAC and snmC-Seq) based on the Allen Reference Atlas. Nissl-stained images show the posterior face of tissue slices (600 µm thickness) used for mouse MOp dissection.

## Supplementary Note

Evaluation of cluster replicability with MetaNeighbor.

**Supplementary Table 1:**

List of datasets, number of cells, and other parameters of each dataset. Data from this study are available via the Neuroscience Multi-omics Archive (NEMO, RRID:SCR_016152) at https://assets.nemoarchive.org/dat-ch1nqb7.

**Supplementary Table 2:**

List of all cells with cluster assignments from 3 computational methods (RNA consensus, SingleCellFusion, LIGER).

**Supplementary Table 3:**

Cluster annotations and unique accession IDs.

**Supplementary Table 4:**

Cluster analysis and metadata for each dataset on its own. Eight individual files:

1. S4a - scRNA SMART
2. S4b - scRNA 10x v3 A
3. S4c - scRNA 10x v2 A
4. S4d - snRNA SMART
5. S4e - snRNA 10x v3 B
6. S4f - snRNA 10x v3 A
7. S4g - Open chromatin (ATAC-seq)
8. S4h - DNA methylation (snmC-seq2)

**Supplementary Table 5:**

Full gene-by-cluster tables for each dataset. Eight individual files:

9. S5a - scRNA SMART
10. S5b - scRNA 10x v3 A
11. S5c - scRNA 10x v2 A
12. S5d - snRNA SMART
13. S5e - snRNA 10x v3 B
14. S5f - snRNA 10x v3 A
15. S5g - Open chromatin (ATAC-seq)
16. S5h - DNA methylation (snmC-seq2)

**Supplementary Table 6:**

For each of the 117 consensus transcriptomic cell types, we performed differential expression (DE) analysis with respect to each of the other cell types. The table reports the top 50 conserved DE genes in each direction for each comparison. Conserved DE genes are significant in at least one dataset, while also having more than two-fold change in the same direction in all but one datasets.

**Supplementary Table 7:**

Enhancers predicted for each cell type based on integrated DNA methylation and ATAC-Seq data using REPTILE.

**Supplementary Table 8:**

List of SingleCellFusion clusters at three levels of cluster resolutions (L0, L1, L2).

## Methods

### Tissue collection and isolation of cells or nuclei (RNA-Seq at Allen Institute, applies to all datasets except snRNA 10x v3 B; Allen)

*Mouse breeding and husbandry:* All procedures were carried out in accordance with Institutional Animal Care and Use Committee protocols at the Allen Institute for Brain Science. Mice were provided food and water *ad libitum* and were maintained on a regular 12-h day/night cycle at no more than five adult animals per cage. For this study, we enriched for neurons by using *Snap25-IRES2-Cre* mice^58^ (MGI:J:220523) crossed to *Ai14*^59^ (MGI: J:220523), which were maintained on the C57BL/6J background (RRID:IMSR_JAX:000664). Animals were euthanized at 53−59 days of postnatal age. Tissue was collected from both males and females (scRNA SMART, snRNA SMART, scRNA 10x v3 A, snRNA 10x v2 A), only males (scRNA 10x v2 A) or only females (snRNA 10x v3 A).

*Single-cell isolation:* We isolated single cells by adapting previously described procedures^5,60^. The brain was dissected, submerged in ACSF^5^, embedded in 2% agarose, and sliced into 250-μm (SMART-Seq) or 350-μm (10x Genomics) coronal sections on a compresstome (Precisionary Instruments). The Allen Mouse Brain Common Coordinate Framework version 3 (CCFv3, RRID:SCR_002978) ^61^ ontology was used to define MOp for dissections.

For SMART-Seq, MOp was microdissected from the slices and dissociated into single cells with 1 mg/ml pronase (Sigma P6911-1G) and processed as previously described^5^. For 10x Genomics, tissue pieces were digested with 30 U/ml papain (Worthington PAP2) in ACSF for 30 mins at 30 °C. Enzymatic digestion was quenched by exchanging the papain solution three times with quenching buffer (ACSF with 1% FBS and 0.2% BSA). The tissue pieces in the quenching buffer were triturated through a fire-polished pipette with 600-µm diameter opening approximately 20 times. The solution was allowed to settle and supernatant containing single cells was transferred to a new tube. Fresh quenching buffer was added to the settled tissue pieces, and trituration and supernatant transfer were repeated using 300-µm and 150-µm fire polished pipettes. The single cell suspension was passed through a 70-µm filter into a 15-ml conical tube with 500 ul of high BSA buffer (ACSF with 1% FBS and 1% BSA) at the bottom to help cushion the cells during centrifugation at 100xg in a swinging bucket centrifuge for 10 minutes. The supernatant was discarded, and the cell pellet was resuspended in quenching buffer.

All cells were collected by fluorescence-activated cell sorting (FACS, BD Aria II, RRID: SCR_018091) using a 130-μm nozzle. Cells were prepared for sorting by passing the suspension through a 70-µm filter and adding DAPI (to the final concentration of 2 ng/ml). Sorting strategy was as previously described^5^, with most cells collected using the tdTomato-positive label. For SMART-Seq, single cells were sorted into individual wells of 8-well PCR strips containing lysis buffer from the SMART-Seq v4 Ultra Low Input RNA Kit for Sequencing (Takara 634894) with RNase inhibitor (0.17 U/μl), immediately frozen on dry ice, and stored at −80 °C. For 10x Genomics, 30,000 cells were sorted within 10 minutes into a tube containing 500 µl of quenching buffer. Each aliquot of 30,000 sorted cells was gently layered on top of 200 µl of high BSA buffer and immediately centrifuged at 230xg for 10 minutes in a swinging bucket centrifuge. Supernatant was removed and 35 µl of buffer was left behind, in which the cell pellet was resuspended. The cell concentration was quantified, and immediately loaded onto the 10x Genomics Chromium controller.

### Tissue collection & nuclei isolation (RNA-Seq at Broad Institute, applies to snRNA 10x v3 B)

*Animal housing:* Animals were group housed with a 12-hour light-dark schedule and allowed to acclimate to their housing environment for two weeks post arrival. All procedures involving animals at MIT were conducted in accordance with the US National Institutes of Health Guide for the Care and Use of Laboratory Animals under protocol number 1115-111-18 and approved by the Massachusetts Institute of Technology Committee on Animal Care. All procedures involving animals at the Broad Institute were conducted in accordance with the US National Institutes of Health Guide for the Care and Use of Laboratory Animals under protocol number 0120-09-16. Samples were collected from both male and female mice.

*Brain preparation prior to 10x nuclei sequencing:* At 60 days of age, C57BL/6J mice were anesthetized by administration of isoflurane in a gas chamber flowing 3% isoflurane for 1 minute. Anesthesia was confirmed by checking for a negative tail pinch response. Animals were moved to a dissection tray and anesthesia was prolonged via a nose cone flowing 3% isoflurane for the duration of the procedure. Transcardial perfusions were performed with ice cold pH 7.4 HEPES buffer containing 110 mM NaCl, 10 mM HEPES, 25 mM glucose, 75 mM sucrose, 7.5 mM MgCl_2_, and 2.5 mM KCl to remove blood from brain and other organs sampled. The brain was removed immediately and frozen for 3 minutes in liquid nitrogen vapor and moved to -80°C for long term storage. A detailed protocol is available at protocols.io^21^.

*Generation of MOp nuclei profiles:* Frozen mouse brains were securely mounted by the cerebellum onto cryostat chucks with OCT embedding compound such that the entire anterior half including the primary motor cortex (MOp) was left exposed and thermally unperturbed. Dissection of 500 µm anterior-posterior (A-P) spans of the MOp was performed by hand in the cryostat using an ophthalmic microscalpel (Feather safety Razor #P-715) precooled to -20°C and donning 4x surgical loupes. Each excised tissue dissectate was placed into a pre-cooled 0.25 ml PCR tube using pre-cooled forceps and stored at -80°C. In order to assess dissection accuracy, 10 µm coronal sections were taken at each 500 µm A-P dissection junction and imaged following Nissl staining. Nuclei were extracted from these frozen tissue dissectates using gentle, detergent-based dissociation, according to a protocol (available at protocols.io) adapted from one generously provided by the McCarroll lab, and loaded into the 10x Chromium v3 system. Reverse transcription and library generation were performed according to the manufacturer’s protocol.

### Epigenomic samples (snATAC-Seq, snmC-Seq2; Salk Institute and UCSD)

*Tissue preparation for nuclei production:* Adult C57BL/6J male mice were purchased from Jackson Laboratories. Brains were extracted from 56-63 day old mice and immediately sectioned into 0.6 mm coronal sections, starting at the frontal pole, in ice-cold dissection media^16^. The primary motor cortex (MOp) was dissected from slices 2 through 5 along the anterior-posterior axis according to the Allen Brain reference Atlas (Extended Data Figure 5). Slices were kept in ice-cold dissection media during dissection and immediately frozen in dry ice for subsequent pooling and nuclei production. For nuclei isolation, the MOp dissected regions from 15-23 animals were pooled for each biological replicate, and two replicates were processed for each region. Nuclei were isolated by flow cytometry as described in previous studies^14,16^. Briefly, nuclei were produced by homogenization in sucrose buffer as described^16^, and the nuclei pellet produced was divided into two aliquots. One aliquot underwent sucrose gradient purification and NeuN labeling (snmC-Seq), and the second went directly to tagmentation (snATAC-seq).

*Bisulfite conversion and library preparation for snmC-Seq2:* Detailed methods for bisulfite conversion and library preparation are previously described for snmC-Seq2^20^, and the protocol is available on protocols.io^35^. The snmC-Seq2 libraries were sequenced using an Illumina Novaseq 6000 instrument (RRID:SCR_016387) with S4 flowcells and 150 bp paired-end mode.

*snATAC-seq data generation:* Combinatorial barcoding single nucleus ATAC-seq was performed as described previously^36,62^. Isolated brain nuclei were pelleted with a swinging bucket centrifuge (500 x g, 5 min, 4°C; 5920R, Eppendorf). Nuclei pellets were resuspended in 1 ml nuclei permeabilization buffer (5 % BSA, 0.2 % IGEPAL-CA630, 1mM DTT and cOmpleteTM, EDTA-free protease inhibitor cocktail (Roche) in PBS) and pelleted again (500 x g, 5 min, 4°C; 5920R, Eppendorf, RRID:SCR_018092). Nuclei were resuspended in 500 µL high salt tagmentation buffer (36.3 mM Tris-acetate (pH = 7.8), 72.6 mM potassium-acetate, 11 mM Mg-acetate, 17.6% DMF) and counted using a hemocytometer. Concentration was adjusted to 4500 nuclei/9 µl, and 4,500 nuclei were dispensed into each well of a 96-well plate. For tagmentation, 1 μL barcoded Tn5 transposomes^62^ were added using a BenchSmart™ 96 (Mettler Toledo, RRID:SCR_018093), mixed five times and incubated for 60 min at 37 °C with shaking (500 rpm). To inhibit the Tn5 reaction, 10 µL of 40 mM EDTA were added to each well with a BenchSmart™ 96 (Mettler Toledo) and the plate was incubated at 37 °C for 15 min with shaking (500 rpm). Next, 20 µL 2 x sort buffer (2 % BSA, 2 mM EDTA in PBS) were added using a BenchSmart™ 96 (Mettler Toledo). All wells were combined into a FACS tube and stained with 3 µM Draq7 (Cell Signaling). Using a SH800 (Sony), 40 nuclei were sorted per well into eight 96-well plates (total of 768 wells) containing 10.5 µL EB (25 pmol primer i7, 25 pmol primer i5, 200 ng BSA (Sigma)). Preparation of sort plates and all downstream pipetting steps were performed on a Biomek i7 Automated Workstation (Beckman Coulter, RRID:SCR_018094). After addition of 1 µL 0.2% SDS, samples were incubated at 55 °C for 7 min with shaking (500 rpm). 1 µL 12.5% Triton-X was added to each well to quench the SDS. Next, 12.5 µL NEBNext High-Fidelity 2× PCR Master Mix (NEB) were added and samples were PCR-amplified (72 °C 5 min, 98 °C 30 s, (98 °C 10 s, 63 °C 30 s, 72°C 60 s) × 12 cycles, held at 12 °C). After PCR, all wells were combined. Libraries were purified according to the MinElute PCR Purification Kit manual (Qiagen) using a vacuum manifold (QIAvac 24 plus, Qiagen) and size selection was performed with SPRI Beads (Beckmann Coulter, 0.55x and 1.5x). Libraries were purified one more time with SPRI Beads (Beckmann Coulter, 1.5x). Libraries were quantified using a Qubit fluorimeter (Life technologies, RRID:SCR_018095) and the nucleosomal pattern was verified using a Tapestation (High Sensitivity D1000, Agilent). The library was sequenced on a HiSeq2500 sequencer (Illumina, RRID:SCR_016383) using custom sequencing primers, 25% spike-in library and following read lengths: 50 + 43 + 37 + 50 (Read1 + Index1 + Index2 + Read2)^18^.

### Genomic library preparation, sequencing and data processing

#### Single cell and single nucleus RNA-Seq (Allen Institute)

For SMART-Seq processing, we performed the procedures with positive and negative controls as previously described^5^. The SMART-Seq v4 (SSv4) Ultra Low Input RNA Kit for Sequencing (Takara Cat# 634894) was used to reverse transcribe poly(A) RNA and amplify full-length cDNA. Samples were amplified for 18 cycles in 8-well strips, in sets of 12–24 strips at a time. All samples proceeded through Nextera XT DNA Library Preparation (Illumina Cat# FC-131-1096) using Nextera XT Index Kit V2 (Illumina Cat# FC-131-2001) and a custom index set (Integrated DNA Technolgies). Nextera XT DNA Library prep was performed according to manufacturer’s instructions, with a modification to reduce the volumes of all reagents and cDNA input to 0.4x or 0.5x of the original protocol.

For 10x v2 processing, we used Chromium Single Cell 3’ Reagent Kit v2 (10x Genomics Cat# 120237). We followed the manufacturer’s instructions for cell capture, barcoding, reverse transcription, cDNA amplification, and library construction. We targeted sequencing depth of 60,000 reads per cell.

For 10x v3 processing, we used the Chromium Single Cell 3’ Reagent Kit v3 (10x Genomics Cat# 1000075). We followed the manufacturer’s instructions for cell capture, barcoding, reverse transcription, cDNA amplification, and library construction. We targeted sequencing depth of 120,000 reads per cell.

#### RNA-Seq data processing and QC (Allen)

Processing of SMART-Seq v4 libraries was performed as described previously^5^. Briefly, libraries were sequenced on an Illumina HiSeq2500 platform (paired-end with read lengths of 50 bp) and Illumina sequencing reads were aligned to GRCm38.p3 (mm10) using a RefSeq annotation gff file retrieved from NCBI on 18 January 2016 (https://www.ncbi.nlm.nih.gov/genome/annotation_euk/all/). Sequence alignment was performed using STAR v2.5.3 ^63^. PCR duplicates were masked and removed using STAR option ‘bamRemoveDuplicates’. Only uniquely aligned reads were used for gene quantification. Gene counts were computed using the R GenomicAlignments package (RRID:SCR_018096)^64^ and summarizeOverlaps function in ‘IntersectionNotEmpty’ mode for exonic and intronic regions separately. For the SSv4 dataset, we only used exonic regions for gene quantification. Cells that met any one of the following criteria were removed: < 100,000 total reads, < 1,000 detected genes (CPM > 0), < 75% of reads aligned to genome, or CG dinucleotide odds ratio > 0.5. Cells were classified into broad classes of excitatory, inhibitory, and non-neuronal based on known markers, and cells with ambiguous identities were removed as doublets^5^.

10x v2 and 10x v3 libraries were sequenced on Illumina NovaSeq 6000 (RRID:SCR_016387) and sequencing reads were aligned to the mouse pre-mRNA reference transcriptome (mm10) using the 10x Genomics CellRanger pipeline (version 3.0.0, RRID:SCR_017344) with default parameters. Cells were classified into broad classes of excitatory, inhibitory, and non-neuronal based on known markers. Low quality cells that fit the following criteria were filtered from clustering analysis. Different filtering criteria for neurons and non-neurons were used as neurons are bigger than non-neuronal cells and contain much more transcripts. For scRNA datasets, neurons with fewer than 2000 detected genes and non-neuronal cells with fewer than 1000 detected genes; for snRNA datasets, neurons with fewer than 1000 detected genes and non-neuronal cells with fewer than 500 detected genes.Doublets were identified using a modified version of the DoubletFinder algorithm^65^ and removed when doublet score > 0.3.

#### Chromatin accessibility (snATAC-Seq) data pre-processing (UCSD)

Paired-end sequencing reads are demultiplexed and then aligned to mm10 reference genome using bwa^66^. After alignment, we converted paired-end reads into fragments and for each fragment, we check the following attributes: 1) mapping quality score MAPQ; 2) whether two ends are appropriately paired according to the alignment flag information; 3) fragment length. We only keep the properly paired fragments whose MAPQ (--min-mapq) is greater than 30 with fragment length less than 1000bp (--max-flen). Because the reads have been sorted based on the names, fragments belonging to the same cell (or barcode) are naturally grouped together which allows for removing PCR duplicates. After alignment and filtration, we used Snaptools (https://github.com/r3fang/SnapTools, RRID:SCR_018097) to generate a snap-format file that contains metadata, cell-by-bin count matrices of a variety of resolutions, cell-by-peak count matrix.

##### Filtering cells by TSS enrichment and unique fragments

The method for calculating enrichment at TSS was adapted from a previously described method ^67^. TSS positions were obtained from the GENCODE database (RRID:SCR_014966). Briefly, Tn5 corrected insertions were aggregated +/-2,000 bp relative (TSS strand-corrected) to each unique TSS genome wide. Then this profile was normalized to the mean accessibility +/-1,900-2,000 bp from the TSS and smoothed every 11bp. The max of the smoothed profile was taken as the TSS enrichment. We then filtered all single cells that had at least 1,000 unique fragments and a TSS enrichment of 10 for all sample sets.

##### Doublet removal

After filtering out low-quality nuclei, we adopt a recently reported algorithm Scrublet (RRID:SCR_018098)^68^ to remove potential doublets for every sample set. Cell-by-peak count matrix are used as input, with default parameters.

##### Clustering

We used the snapATAC pipeline^62^ to identify cell clusters with binarized cell-by-bin matrix in 5kb resolution as the input. Cell clusters were annotated to cell type by checking chromatin accessibility along the body of marker genes. Then another round of clustering were performed on MGE- and CGE-derived inhibitory GABA-ergic interneurons, in order to identify sub-cell types.

#### DNA methylation (snmC-Seq) data pre-processing (Salk)

##### Mapping and feature count pipeline for snmC-Seq2

We implemented a versatile mapping pipeline (cemba-data.rtfd.io) for all the single-cell methylome based technologies developed by our group^16,20,37^. The main steps of this pipeline included: 1) Demultiplexing FASTQ files into single-cell files; 2) Reads level QC; 3) Mapping; 4) BAM file processing and QC; 5) final molecular profile generation. The details of the five steps for snmC-seq2 were described previously^69^. We mapped all the reads onto the mouse mm10 genome. After mapping, we calculated the methyl-cytosine counts and total cytosine counts in two sets of genome regions for each cell: the non-overlapping 100kb bins tiling the mm10 genome, which was used for methylation-based clustering analysis, and gene body regions ± 2kb, which is used for cluster annotation and cross modality integration.

##### Quality control and cell filtering

We filtered the cells based on these main quality metrics: 1) The rate of bisulfite non-conversion as estimated by the rate of methylation at CCC positions (mCCC) < 0.03. mCCC rate reliably estimates the upper bound of bisulfite non-conversion rate^16^, 2) overall mCG rate > 0.5, 3) overall mCH rate < 0.2, 4) total final reads (combining R1 and R2) > 500,000, 5) Total mapping rate (using Bismark^70^) > 0.5.

##### Preprocessing and clustering

The clustering steps of snmC-seq2 data were described previously^37^. In brief, we calculated posterior mCH and mCG rate based on beta-binomial distribution for the non-overlapping 100kb bins matrix, we then selected top 3000 highly variable features to perform PCA and find dominant PCs for mCH and mCG separately. We concatenate PCs from both methylation types together to construct a KNN graph, and ran the Leiden community detection algorithm^71^ repeatedly to get the consensus clustering results. The stopping criteria of clustering considered number of marker genes, accuracy of the reproducible supervised model based on the cluster assignments, and minimum cluster size. We performed the clustering in two iterations to get major types and fine-grained types for comparison with other modalities in further integration.

### Computational Analysis

#### Transcriptome analysis (Fig. 2)

##### Clustering individual datasets

Clustering for each sc/snRNASeq dataset was performed independently using the R package *scrattch.hicat*^5^ (RRID:SCR_018099, available at https://github.com/AllenInstitute/scrattch.hicat). In addition to classical single-cell clustering processing steps provided by other tools such as Seurat, this package supports iterative clustering by making successively finer splits while ensuring all pairs of clusters, even at the finest level, are separable by stringent differential gene expression criteria^5^. For the scRNA 10x datasets, we used q1.th = 0.4, q.diff.th=0.7, de.score.th=150, min.cells=10. For the snRNA 10x datasets, we used q1.th=0.3, q.diff.th=0.7, de.score.th=100, min.cells=10. For the scRNA SMART datasets, we used q1.th = 0.5, q.diff.th=0.7, de.score.th=150, min.cells=4. For the snRNA SMART dataset, we used q1.th=0.4, q.diff.th=0.7, de.score.th=100, min.cells=4. We further performed consensus clustering by repeating iterative clustering on a subsample of 80% of cells, resampled 100 times, followed by final clustering based on the co-clustering probability matrix. Using this procedure, we could fine tune cluster boundaries as well as assess cluster uncertainty.

##### Joint clustering of multiple datasets

To provide a consensus cell type taxonomy across all transcriptomic datasets, we developed a novel integrative clustering analysis across multiple data modalities. This procedure is available via the *harmonize* function of the *scrattch.hicat* package. Unlike Seurat/CCA^40^, which aim to find aligned common reduced dimensions across multiple datasets, this method directly builds a common adjacency graph using the cells from all datasets, then applies the Louvain community detection algorithm^72^. We extended the cluster merging algorithm in the *scrattch.hicat* package to ensure that all clusters can be separated by conserved DE genes across platforms. The *i_harmonize* function, similar to the *iter_clust* function in the single dataset clustering pipeline, applies integrative clustering across datasets iteratively while ensuring all the clusters at each iteration are separable by conserved DE genes.

To build a common adjacency matrix incorporating samples from all the datasets, we first chose a subset of datasets which we used as “reference datasets.” Reference datasets provide the most sensitive gene detection and/or comprehensive cell type coverage. For this study, we used 10x v2 single cell dataset from Allen (scRNA 10x v2 A) and 10x v3 single nucleus dataset from Broad (snRNA 10x v3 B) as references, as both are large datasets that provide comprehensive cell type coverage and relatively sensitive gene detection.

The key steps of the pipeline are outlined below:

1. **Perform single-dataset clustering** (Methods described above).
2. **Select anchor cells for each reference dataset.** For each reference dataset (scRNA 10x v2 A or snRNA 10x v3 B), we randomly sampled up to 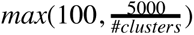 anchor cells per cluster to normalize coverage for each cell type. This is the only step that uses the dataset-specific clustering information.
3. **Select highly variable genes (HVG).** Highly variable gene selection and dimensional reduction by principal components analysis (PCA) were performed using the *scrattch.hicat* package. We removed PCs with a Pearson correlation coefficient of more than 0.7 with log2(Ngenes). This step was implemented to mitigate the effect of cell/nucleus quality on gene expression variability, and to select only biologically relevant PCs. For each remaining PC, Z scores were calculated for gene loadings. The top 100 genes with absolute Z score greater than 2 were selected as HVGs. The HVGs from each reference dataset were combined.
4. **Compute K nearest neighbors (KNN).** For each cell in each query dataset, we computed its K nearest neighbors (k=15) among anchor cells in each reference dataset (scRNA 10x v2 A or snRNA 10x v3 B), based on the highly variable genes selected above. The RANN package was used to compute KNN based on the Euclidean distance when the query and reference dataset is the same. To compute nearest neighbors across datasets, we used correlation as a similarity metric.
5. **Compute the Jaccard similarity.** For every pair of cells from all datasets, we compute their Jaccard similarity, defined as the ratio of the number of shared K nearest neighbors (among all anchors cells from all the reference datasets) divided by the number of combined K nearest neighbors.
6. **Perform Louvain clustering.**
7. **Merge clusters.** To ensure that every pair of clusters are separable by conserved differentially expressed (DE) genes across all datasets, for each cluster, we first identified the top 3 most similar clusters. For each pair of such closely-related clusters, we computed the differentially expressed genes in each dataset. We focus on the conserved DE genes that are significant in at least one dataset, while also having more than two-fold change in the same direction in all but one datasets. We then compute the overall statistical significance based on such conserved DE genes for each dataset independently. If any of the datasets pass our DE gene criteria described in the “clustering” section, the pair of clusters remain separated; otherwise they are merged. DE genes were recomputed for the merged clusters, and the process was repeated until all clusters are separable by the conserved DE genes criteria. If one cluster has fewer than the minimal number of cells in a dataset (4 cells for SMART-Seq and 10 cells for 10x), then this dataset is not used for DE gene computation for all pairs involving the given cluster. This step allows detection of unique clusters absent in some platforms.
8. **Iterative clustering.** Repeat step 1-6 for cells within each cluster to gain finer resolution clusters until no more clusters can be found.
9. **Final compilation and merging of clusters.** Concatenate all the clusters from all the iterative clustering steps, and perform final merging as described in step 6.

##### Marker gene selection

For each pair of clusters, we computed the conserved DE genes, i.e. those which are significantly DE in one at least dataset, with ≥2-fold change in expression in the same direction among 70% of datasets. To allow computation of DE genes involving cell types only present in a subset of datasets, only the datasets with enough cells (based on min.cells parameter) for both cell types under comparison were used for DE gene calculation. We selected the top 50 genes in each direction. After pooling genes from all pairwise comparisons, we identified a total of 3,792 marker genes (Supplementary Table 6).

##### Imputation

To facilitate direct comparison, we projected gene expression of all datasets to the space of a given reference dataset. To do that, we leveraged the KNN matrices computed during the iterative joint clustering step to adjust the expression values for systematic differences between datasets. During each iteration of the joint clustering, for cells in each dataset, we used the average gene expression of their k nearest neighbors among the anchor cells from the reference dataset as the adjusted expression in the reference space. At the top-level clustering, we imputed the expression for all genes. For each subsequent iteration, we only imputed the expression of the high-variance genes and the conserved DE genes for the clusters defined in that iteration. We used this iterative approach for imputation because the nearest neighbors based on the genes chosen at the top level may not reflect the distinction between the finer types, and the imputed values for the DE genes that define the finer types consequently are not accurate based on these nearest neighbors. Therefore, we deferred imputation of the DE genes between the finer types to the iteration when these types were defined. This method is provided in the **impute_knn_global** function in scrattch.hicat packaget^5^. We imputed the gene expression matrix for both reference datasets used in the integrative clustering.

##### Building a cell-type taxonomy tree

We first compute the average of the adjusted expression of marker genes for each cluster. This average was computed using each of the two reference datasets (scRNA 10x v2 A, snRNA 10x v3 B). Then, the two matrices were concatenated. We constructed a hierarchy (tree) using the **build_dend_harmonize** function in *scrattch.hicat* packaget^5^.

##### Dimensionality reduction by UMAP

We performed principal component analysis (PCA) based on imputed gene expression matrices of 3,792 marker genes using 10x single nuclei dataset from Broad as the reference, and selected the top 50 principal components (93% variance explained). We removed PCs with Pearson correlation coefficient >0.6 with the log2(Ngenes) to reduce bias related to the number of detected genes. Uniform Manifold Approximation and Projection (UMAP) was used to embed the cells in two dimensions with parameters nn.neighbors=25 and md=0.3^73^.

#### MetaNeighbor analysis (Fig. 2g,h)

To quantify replicability of clusters across the 7 transcriptomic datasets, we applied a modified version of unsupervised MetaNeighbor (RRID:SCR_016727)^31^. MetaNeighbor uses a neighbor voting algorithm and a cross-dataset validation scheme to quantify cluster similarity across multiple datasets. It requires a set of unnormalized datasets, a set of cluster labels and a set of highly variable genes. We used the raw count data for all cells passing QC criteria for the 7 single cell transcriptome datasets, as well as the labels obtained through independent clustering (Supplementary Table 5). We used MetaNeighbor’s *variableGenes* procedure to select 310 highly variable genes that were detected as highly variable across all datasets.

We defined replicable clusters in a two-step procedure: first we quantified the similarity between clusters across datasets, then we extracted groups of highly similar clusters, or “meta-clusters”. We used the *MetaNeighborUS* function to obtain an initial similarity matrix between clusters. By default, cluster similarity is quantified as a one-vs-all area under the receiver-operator curve (AUROC): given a training cluster (in one dataset), we ask how similar cells from a test cluster (in another dataset) are to training cells, compared to all other cells in the test dataset. To make cluster matching more stringent, we transformed the one-vs-all AUROC matrix into a one-vs-best AUROC matrix: instead of ranking test cells among all cells from the test dataset, we only compare them to cells from the best matching cluster. This modification ensures that only the best match can have an AUROC > 0.5, facilitating identification of reciprocal best hits. For interpretability and computational efficiency, we adopted the following convention: the best matching cluster’s AUROC was obtained by comparing it to the second best matching cluster, the second best cluster’s AUROC was obtained by computing 1-AUROC of the best matching cluster, and all other clusters obtained an AUROC of 0, as we were only interested in finding best matches. To extract meta-clusters, we interpreted the one-vs-best AUROC as a graph where nodes are clusters and edges connect nodes if they are reciprocal best hits. We define meta-clusters as connected components in this graph. We can obtain more robust meta-clusters by requiring that best hits exceed some AUROC threshold. In practice, we noted that one-vs-best AUROC > 0.7 offered a good balance between the number of meta-clusters and reproducibility strength. For scalability, we modified MetaNeighbor in the following ways. In the *MetaNeighborUS* function, we removed the rank standardization of the cell-cell similarity network (by setting parameter *fast_version* to *TRUE*) and the node degree normalization of the neighbor voting, enabling analytical simplifications of the neighbor voting procedure. The *variableGenes* procedure was applied to a random subset of 50,000 cells for datasets exceeding that size.

#### Epigenomic data (Fig. 3)

##### Cluster analysis for snmC-Seq

We concatenate principal components from both methylation types (CG and CH) together, and use these to construct a KNN graph followed by Leiden community detection^71^. We repeat the cluster analysis several times to get consensus clustering results. The stopping criteria of clustering considered number of marker genes, accuracy of the reproducible supervised model based on the cluster assignments, and minimum cluster size. We performed the clustering in two iterations to get major types and fine-grained cell types for comparison with other modalities in further integration.

Two-dimensional embedding using t-distributed stochastic neighbor embedding^74^ (tSNE; perplexity = 30) was calculated based on the top principal components using the implementation from the scanpy package^75^.

#### Multimodality integration (Fig. 4)

##### Computational data integration with LIGER

We used LIGER (RRID:SCR_018100) to integrate the single-cell transcriptomic and epigenomic data as previously described in the LIGER paper^42^, with one modification. We used the *optimizeALS* function in the LIGER package to perform joint factorization on all datasets except methylation (7 RNA datasets and one ATAC dataset) to infer shared (*W*) and dataset-specific (*V_i_*) metagene factors and cell factor loadings (*H_i_*). We then used the resulting *W* to calculate cell factor loadings (*H_i_*) for the methylation data using the *solveNNLS* function in the LIGER package. We found that this strategy yielded better integration than jointly factorizing all 8 datasets, possibly because the inverse relationship and massive dataset size imbalance between methylation and all other datasets complicates the learning of shared metagenes. Our analysis used only the cells annotated by each data-generating group as passing quality control. We did not perform any data imputation or smoothing, but simply normalized and scaled the raw cell-by-gene count matrices from each dataset using the *normalize* and *scaleNotCenter* functions in the LIGER package. We next used the *quantileAlignSNF* function with default settings to perform quantile normalization of cell factor matrices (*H_i_*) from all 8 datasets. Finally, we performed Louvain clustering on the normalized cell factor matrices (*H_i_*) to obtain joint clusters. We performed two rounds of integration and joint clustering; in the first round, we separately integrated all neurons across datasets and all glia across datasets. We then performed a second round of integration and clustering separately for each of four neuronal subclasses: excitatory intratelencephalic (IT) neurons, excitatory non-IT neurons, medial ganglionic eminence (MGE) interneurons, and caudal ganglionic eminence (CGE) interneurons. We used *k*=40 factors for the non-neuron analysis, *k*=30 for the first-round neuron analysis, and *k*=20 for all of the second-round analyses.

##### Computational integration with SingleCellFusion

SingleCellFusion^37^ is designed to robustly integrate DNA methylation, ATAC-Seq and/or RNA-Seq data. We applied SingleCellFusion iteratively to integrate all neurons from 8 datasets (Supplementary Table 1) and jointly call cell clusters. To integrate both the broad and fine-grained cell types, we performed 3 rounds of integration. For every cell cluster generated in the previous round, it is further split into smaller clusters by re-applying SCF on cells in that cluster only. In the first round, we run SCF on all neurons from 8 datasets and get 10 broad neuronal clusters. Rounds 2 and 3 generate 29 clusters and 56 more fine-grained clusters, respectively (Supplementary Table 3).

The procedure comprises 4 major steps: preprocessing: within-modality smoothing, cross-modality imputation, and clustering and visualization.

1. **Preprocessing.** We define a gene-by-cell feature matrix for each dataset. Droplet-based RNA-seq features (10x) are log_10_(CPM+1) normalized; Full-length RNA-seq (SMART-seq) features are log_10_(TPM+1) normalized. snATAC-seq data is represented by read counts within gene body, normalized by log_10_(RPM+1), where CPM stands for counts per million reads mapped (counts normalized), TPM stands for transcripts per million reads mapped (length normalized), and RPM stands for reads per million reads mapped (length normalized), respectively. DNA methylation data is represented by the mean gene body mCH level, normalized by the global (genome-wide) mean mCH level for each cell. For each dataset, we only used high-quality cells (passed QC) and highly variable genes (n=4,000∼6,300) for further analysis. To select highly variable genes, for RNA-seq and ATAC-seq datasets, we first remove genes that are expressed in < 1% of cells. We then divide the remaining genes into 10 bins according to their mean expression across cells (CPM). For each bin, except for the one with the most expressions, we select top 30% of genes with the most expression dispersion (variance/mean) as the highly variable genes. For the DNA methylation dataset, we first select genes that have > 20 cytosine coverage in more than 95% of cells, then divide the remaining genes into 10 bins according to their mean normalized mCH level--raw mCH level normalized by the global mCH for each cell. For each bin, we select top 30% of genes with the most variance as the highly variable genes.
2. **Within-modality smoothing.** To reduce the sparsity and noise of feature matrices, we share information among cells with similar profiles using data diffusion. The procedure is adapted from ^76^ and described in detail in ^37^. Here we exactly followed (Luo et al 2019 bioRxiv)^37^ with [ndim=50, k=30, ka=5] for all datasets, and [p=0.7] for RNA-seq datasets, [p=0.9] for the DNA methylation dataset, and [p=0.1] for the ATAC-seq dataset.
3. **Cross-modality imputation by Restricted k-Partners (RKP)**. To integrate all 8 datasets, we impute the scRNA_10x_v2_A gene features for cells in all 7 other datasets. The imputation is done in pairwise between the scRNA_10x_v2_A dataset and one other dataset. For each pairwise imputation, we followed the procedure described in ^37^ with 20 RKP and relaxation parameter 3 [k=20, z=3]. Instead of using Euclidean distance in a low-dimensional space, we here use the (flipped) spearman correlation coefficient across genes that are highly variable in both datasets as the distance metric between cells in 2 different modalities.
4. **Clustering and visualization.** We start from a cell-by-feature matrix, where cells include all cells from 8 datasets and features are highly variable genes of the scRNA_10x_v2_A dataset. We reduce the dimensionality of features into top 50 Principal Components. Next, we perform UMAP embedding on the PC matrix [n_neighbors=60, min_dist=0.5]. Finally, we perform Leiden clustering on the kNN graph (symmetrized, unweighted) generated from the final PC matrix [Euclidean distance, k=30, resolution=0.1].

##### Fig. 4 related panel-specific analysis

Figure 4h We created the embedding of the cluster centroids using the imputed scRNA_10x_v2_A gene features (log10(CPM+1)) for all cells from the 8 different datasets generated from SingleCellFusion integration. Clusters are defined by individual dataset clusterings and by the joint clustering with SingleCellFusion. Cluster centroids are calculated by the mean imputed scRNA_10x_v2_A gene profiles across cells. After getting a gene-by-cluster matrix, we apply PCA to reduce to 50 feature dimensions, followed by applying a UMAP embedding with min_dist=0.7 and n_neighbors=10.

Figure 4i To compare molecular signals across data modalities, all signals are normalized to [0, 1]. This is achieved by first getting molecular signals by dataset-specific normalization (Step1), followed by a linear transformation (Step2). In Step1, for SMART-seq datasets, we show log10(TPM+1); for 10x RNA-seq datasets, we show log10(CPM+1); for the ATAC-seq dataset, we show log10(RPM+1) normalized gene body counts, and for DNA methylation we show gene body mCH normalized by global mCH level of each cell. For Step2, we apply a linear transformation to map the range of the signal to [0, 1]. For datasets other than DNA methylation, we apply the following formula:

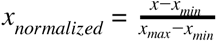

Where *x* is the dataset-specific gene-level signal for a cell, *x_min_* and *x_max_* are defined as the bottom 2 percentile and top 2 percentile of *x* across all cells, respectively. For the DNA methylation dataset, we apply the following formula:

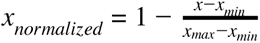

, with which signals are still mapped to [0, 1] but flipped--a high signal on the plot means a low DNA methylation level. We do this to align DNA methylation signals with gene expression (and open chromatin) signals, because DNA methylation is a repressive marker of gene expression and negatively correlates with it. Besides, *x_min_* and *x_max_ x* across all cells, respectively. are defined as the bottom 2 percentile and top 50 percentile of

Figure 4j For each gene, cell-level signals are normalized the same way as described in Step1 of Figure 4i. Cluster level signals are the mean cell-level signals across cells in clusters. After getting gene-by-cluster matrices this way, for non-DNA methylation datasets, the matrices are further normalized by the maximum of each cluster (column); for DNA methylation datasets, no further normalization is done, for they are already normalized by cell.

**Extended Data Figure 5g,h** The heatmaps show pairwise Spearman correlation coefficients between the centroids of cells from each cell type (SingleCellFusion) and each dataset, using the gene expression levels (log_10_(CPM+1); measured or imputed by SingleCellFusion) of the scRNA_10x_v2_A dataset as features. Centroid-level profiles are computed as the average of cell-level profiles across cells from the same cell type and the same dataset. The row and column orderings are the same, generated by a hierarchical clustering on the above defined centroid-level features with average linkage and euclidean distance. 5f shows the correlations between broad-level joint clusterings (10 subclasses; SingleCellFusion L0; Supplementary Table 8); 5g shows those between fine-level joint clusterings (56 clusters in total; not all are shown; SingleCellFusion L2; Supplementary Table 8) for four example broad-level subclasses (MGE, CGE, L2/3 IT, L4/5 IT).

#### Extended Data Figure 5i (Agreement metric)

We calculated dataset agreement metrics as described in the LIGER paper ^42^. Briefly, we performed dimensionality reduction using either NMF (for LIGER) or PCA (for SingleCellFusion) and built a k-nearest neighbor graph for each individual dataset. Then we built a *k*-nearest neighbor graph using the joint latent space from either LIGER or SingleCellFusion and calculated what fraction of the nearest neighbors from individual datasets were still nearest neighbors in the joint space. This metric assesses how well the joint latent space preserves the structure of each individual dataset. An agreement metric close to zero indicates poor preservation of structure from individual datasets, while an agreement metric close to 1 ideally preserves the structure.

#### Extended Data Figure 5j (Alignment metric)

We calculated dataset alignment metrics as described in the LIGER ^42^ and Seurat ^40^ papers, except that we first downsampled cells so that the cluster proportions and total number of cells were identical across all datasets. Then we built a *k*-nearest neighbor graph using the joint latent space from either LIGER or SingleCellFusion and calculated what fraction of the nearest neighbors around each point come from each dataset. We then normalized the metric to be between 0 (no alignment) and 1 (perfect mixing of datasets). This metric assesses how well the joint latent space aligns the datasets. Note that maximizing alignment and maximizing agreement are competing objectives. For example, it is possible to trivially maximize alignment by randomly mixing cells from all datasets according to a spherical Gaussian distribution; conversely, one could trivially maximize agreement by simply assigning non-overlapping latent representations to all datasets. However, methods must balance these competing objectives to score highly on both alignment and agreement metrics.

#### Extended Data Figure 5k

To get cluster-level gene signals, we first get normalized cell-level signals the same way as Step1 of Figure 4i, followed by taking the mean cell-level signals across cells in clusters.

#### Analysis of enhancers (Fig. 5)

##### Epigenome Cluster Level

Based on the cell-cell integration in Figure 4, in order to have enough whole-genome coverage of each cell type, we further merged the co-clusters into a higher level to increase the coverage of each cluster, which we termed as the epigenome cluster level.

##### DMR and Peak Calling

For DMR calling in the snmC-seq2 data, we merged single-cell ALLC files into the pseudo-bulk level for each cluster, and then used methylpy^77^ DMRfind function to calculate mCG DMRs across all clusters. The base call of each paired CpG sites was added up before analysis. In brief, the methylpy function used a permutation-based root-mean-square test of goodness-of-fit to identify differentially methylated sites (DMS) simultaneously across all samples, and then merge the DMS within 250bp into DMR. Hypo-DMR and Hyper-DMR were then assigned to each sample by examining the residue of observed counts from the expected counts. We also filtered the DMRs by requiring the maximin difference of mCG rate between clusters larger than 0.3. For peak calling in the snATAC data, we extracted all the fragments for each cluster, and then performed peak calling on each aggregate profile using MACS2^78^ with parameter: “--nomodel --shift -100 --ext 200 --qval 1e-2 –B --SPMR”. We used the “*bedtools intersect*” with the “-wa -u” parameter *to calculate* DMR and ATAC peak overlaps.

##### Saturation analysis

To investigate the efficiency of regulatory elements identification in terms of cell number in the epigenomic data, we did a saturation analysis using the two most abundant cell types: the L2/3 IT and the L6 CT excitatory neurons, the total reads assigned to these two cell types were comparable to bulk-seq. We subsampled a different number of cells without replacement in each cluster three times when having enough cells, and used cells from each replicate separately when possible. In the last group, we used all the cells for each cell type as a maximum reference. For methylome data, We call DMRs between L2/3 IT and L6 CT within each cell number group. Peaks are called for each cell type group.

##### REPTILE enhancer prediction

We performed enhancer prediction using the REPTILE^79^ algorithm. The REPTILE is a random-forest-based supervised method that incorporates different sources of epigenomic profiles with base-level DNA methylation data to learn and then distinguish the epigenomic signatures of enhancers and genomic background. We trained the model in a similar way as in previous studies^79,80^ using CG methylation, chromatin accessibility of each epigenome clusters and mouse embryonic stem cells (mESC). The model was first trained on mESC data and then predicted a quantitative score we termed enhancer score for each cell type’s DMRs. The positives were 2kb regions centered at the summits of top 5,000 EP300 peaks in mESCs. Negatives include randomly chosen 5,000 promoters and 30,000 2kb genomic bins. The bins have no overlap with any positives or promoters^80^. Methylation and chromatin accessibility profiles in bigwig format for mESC were from the mouse ENCODE project^80^. The mCG rate bigwig file was generated from cell type-merged ALLC files using in-house python script. For chromatin accessibility of each cell type, we merged all fragments from snATAC-seq cells that assigned to this cell type in the integration analysis and used “*deeptools bamcoverage*” to generate CPM normalized bigwig files. All bigwig files’ bin size was 50bp.

##### Motif Enrichment Analysis

We used 724 motif PWMs from the JASPAR 2020 CORE vertebrates database^81^, where each motif was able to assign corresponding mouse transcription factor genes. For each set of REPTILE predicted enhancers, we standardized the region length into center ± 250bp and used the FIMO tool from the MEME suite^82^ to scan the motifs in each enhancer with log odds p-value < 10^-6^ as the threshold of motif hit. To calculate motif enrichment, we use the adult non-neuronal mouse tissue DMRs^46^ as background regions. We subtracted enhancers in the region set from the background, and then scanned the motifs in background regions using the same approach. We then used Fisher’s exact test to find motifs enriched in the region set, and the Benjamini-Hochberg procedure to correct multiple tests. Transcription factors with significant motif enrichment were grouped by TFClass^48^ classification. Genes within the same group share very similar motifs.

#### Cluster validation analysis (Fig. 6)

##### Downsampling analysis of cluster number (Fig. 6a-e)

###### Preprocessing

Preprocessing is done in the same way as described in the section of Computational integration with SingleCellFusion. After preprocessing, we get a gene-by-cell feature matrix for each dataset. Only neuronal cells passing QC (Supplementary Table 1) and highly variable genes for each dataset are included.

###### Clustering (Fig. 6a)

Clustering takes 3 steps. We first reduce feature dimensions by PCA [n=50]. We then build a k-nearest neighbor graph [k=30] between cells using the Euclidean distance in the Principal Component space. We finally apply the Leiden clustering algorithm with a fixed resolution parameter [r=6]. For each dataset, we report the number of clusters as a function of the number of cells randomly downsampled from the full dataset. Error bars show the standard error of the mean of [n=10] repeats of downsampling.

###### Clustering with within-modality cross validation (Fig. 6d)

This analysis aims to estimate the “optimal” number of clusters of a dataset, by testing which clustering granularity best preserves the gene-level features of cells. For a given dataset--a gene-by-cell matrix, we first randomly split gene features into 2 sets, for clustering and validation, respectively. To avoid any potential linkage, the split is done by separating chromosomes into 2 sets, such that genes from the same chromosomes are always in the same set. We then perform Leiden clustering (as described in methods related to Fig. 6a) on all cells using the clustering feature set only with different clustering resolutions. After clustering, every cell in the dataset gets a cluster label. We next randomly separate those cells into 2 sets--for training and testing, respectively. Using training-set cells, we train a supervised model to predict the validation set gene features based on cluster assignments. Assuming a R2 loss, this is equivalent to calculating the cluster centroid of each cluster in the space of validation gene set using training-set cells only. Finally, we apply the model to cells in the test set, and evaluate the mean squared error of model performance. This is equivalent to estimating the mean squared distance between individual cells in the test set to its cluster centroid calculated by training set. As a function of number of clusters (by varying the resolution parameter in Leiden clustering), we observe a U-shaped curve of mean squared error, because both under-splitting and over-splitting results in high mean squared error. The minimum point of the curve represents the most plausible clustering resolution. Applying this scheme to each dataset and different downsampling levels of cells, we report in Fig. 6d the number of clusters as a function of the number of cells, for each dataset. For robustness, random split of gene features are repeated n=5 times; random split of cells are repeated n=5 times with k=5 fold cross validations each time.

###### Clustering with cross-modality cross validation (Fig. 6e)

Extending the within-modality clustering cross validation scheme used in Fig. 6d, we developed a cross-modality cross validation method, by combining the previously described within-dataset cross-validation method with a joint clustering method--SingleCellFusion. First of all, similar to within-dataset cross validation, we first randomly split gene features into clustering and validation set for all datasets. We then generate integrated clusterings across data modalities by applying SingleCellFusion on all cells and half of the gene features (the clustering feature set). After clustering, we estimate the mean squared error of clustering on the validation feature set as described above for each dataset on its own. Applying this scheme to different downsampling levels of cells, we report in Fig. 6e the number of clusters as a function of the number of cells from each dataset.

###### Clustering on network of samples (Conos) analysis (Fig. 6g-i)

To evaluate the extent to which different cell subpopulations were supported by different platforms, we assessed the difference in the ability to recover the corresponding cell with and without within-platform comparisons. The clustering of cells was performed using Conos^55^, using walktrap community detection method to detect hierarchical cell populations. The stability of the resulting hierarchical clustering result was estimated as follows: 20 random cell subsampling rounds were performed, each drawing random 95% of cells from each dataset, repeating the walktrap hierarchical clustering procedure. For each node in the original walktrap tree, we evaluated stability as a minimum of specificity and sensitivity relative to the ensemble of subsampled trees by finding the best matching subtree. To evaluate the ability to recover subpopulations based on cross-platform comparisons only, we removed within-platform edges (those connecting datasets generated by the same platform) in the joint graph (generated by Conos). This way the subpopulation is detected only if it is aggregated based on its mapping to the other platform. The modified approach will facilitate the grouping of cell population that are common in the different platforms as it removed the platform-specific information in the joint graph.

To assess similarity of expression profiles detected by different platforms for a given cell type (Fig. 6i), we used Jensen-Shannon divergence to assess the overall similarity of gene expression patterns between the four RNA-Seq platforms (scRNA 10x v3 A, snRNA 10x v3 B, scRNA SMART and snRNA SMART). Specifically, 1000 cells were sampled from each cell type for each platform. If the number of cells from a cell type is smaller than 1000 cells, sampling with replacement was performed. Cell types that accounted for less than 1% (<300 cells) in any specific platform were omitted. The molecules detected for each gene were then aggregated across all sampled cells for each cell type in each platform. The counts were normalized by the total number of molecules for each cell type / platform, and Jensen-Shannon divergence was calculated.

#### Integrated analyses: trade-off between replicability and resolution and cluster consistency (Fig. 6f, j)

We collected the clusters obtained with the 4 integrative clustering methods described previously (Conos, LIGER, RNA consensus clustering from Figure 2, SingleCellFusion), as well as the “subclass” level from the independent clustering of the RNA datasets. Each integrative method returned clusters at two granularity levels, we named the coarser level of clustering L1 and the finer level of clustering L2 clusters. We focused our analyses on the neuron clusters of the transcriptomic data, as we wished to investigate the agreement of neuron cluster hierarchies.

To quantify replicability, we used the same modified version of MetaNeighbor, same datasets and same variable genes as defined above (see “MetaNeighbor analysis”). We used the one-vs-best AUROC to obtain cluster similarity scores, then computed an average AUROC score per integrated cluster (averaged over every pair of datasets in which the cluster is present). For every method, we reported the median AUROC across integrated clusters as the final reproducibility score. To quantify the overall similarity of the clustering results, we computed the Adjusted Rand Index (ARI). When necessary, we restricted the ARI computation to the intersection of labeled cells (the intersection being recomputed for every pair of methods).

## References

1. Zeng, H. & Sanes, J. R. Neuronal cell-type classification: challenges, opportunities and the path forward. Nat. Rev. Neurosci. 18, 530–546 (2017).

2. Ramon y Cajal, S. Histologie du système nerveux de l’homme et des vertébrés. *Maloine*, Paris 2, 153–173 (1911).

3. Zeisel, A. et al. Molecular Architecture of the Mouse Nervous System. Cell 174, 999–1014.e22 (2018).

4. Saunders, A. et al. Molecular Diversity and Specializations among the Cells of the Adult Mouse Brain. Cell 174, 1015–1030.e16 (2018).

5. Tasic, B. et al. Shared and distinct transcriptomic cell types across neocortical areas. Nature 563, 72–78 (2018).

6. Cadwell, C. R., Bhaduri, A., Mostajo-Radji, M. A., Keefe, M. G. & Nowakowski, T. J. Development and Arealization of the Cerebral Cortex. Neuron 103, 980–1004 (2019).

7. Telley, L. et al. Temporal patterning of apical progenitors and their daughter neurons in the developing neocortex. Science 364, (2019).

8. Fishell, G. & Kepecs, A. Interneuron Types as Attractors and Controllers. Annu. Rev. Neurosci. (2019) doi:10.1146/annurev-neuro-070918-050421.

9. Mukamel, E. A. & Ngai, J. Perspectives on defining cell types in the brain. Curr. Opin. Neurobiol. 56, 61–68 (2018).

10. Waddington, C. H. The strategy of the genes. (Routledge, 1957).

11. Paul, A. et al. Transcriptional Architecture of Synaptic Communication Delineates GABAergic Neuron Identity. Cell 171, 522–539.e20 (2017).

12. Mure, L. S. et al. Diurnal transcriptome atlas of a primate across major neural and peripheral tissues. Science 359, (2018).

13. Lacar, B. et al. Nuclear RNA-seq of single neurons reveals molecular signatures of activation. Nat. Commun. 7, 11022 (2016).

14. Lister, R. et al. Global epigenomic reconfiguration during mammalian brain development. Science 341, 1237905–1237905 (2013).

15. Price, A. J. et al. Divergent neuronal DNA methylation patterns across human cortical development reveal critical periods and a unique role of CpH methylation. Genome Biol. 20, 196 (2019).

16. Luo, C. et al. Single-cell methylomes identify neuronal subtypes and regulatory elements in mammalian cortex. Science 357, 600–604 (2017).

17. Mo, A. et al. Epigenomic Signatures of Neuronal Diversity in the Mammalian Brain. Neuron 86, 1369–1384 (2015).

18. Preissl, S. et al. Single-nucleus analysis of accessible chromatin in developing mouse forebrain reveals cell-type-specific transcriptional regulation. Nat. Neurosci. 21, 432–439 (2018).

19. Ecker, J. R. et al. The BRAIN Initiative Cell Census Consortium: Lessons Learned toward Generating a Comprehensive Brain Cell Atlas. Neuron 96, 542–557 (2017).

20. Luo, C. et al. Robust single-cell DNA methylome profiling with snmC-seq2. Nat. Commun. 9, 3824 (2018).

21. Vanderburg, C. et al. Fresh Frozen Mouse Brain Preparation (for Single Nuclei Sequencing). (2020) doi:10.17504/protocols.io.bcbrism6.

22. Hodge, R. D. et al. Conserved cell types with divergent features in human versus mouse cortex. Nature 573, 61–68 (2019).

23. Cell Type Nomenclature - brain-map.org. https://portal.brain-map.org/explore/classes/nomenclature.

24. Economo, M. N. et al. Distinct descending motor cortex pathways and their roles in movement. Nature 563, 79–84 (2018).

25. Brodmann, K. Brodmann’s: Localisation in the Cerebral Cortex. (Springer Science & Business Media, 2007).

26. Yamawaki, N., Borges, K., Suter, B. A., Harris, K. D. & Shepherd, G. M. G. A genuine layer 4 in motor cortex with prototypical synaptic circuit connectivity. Elife 3, e05422 (2014).

27. Jabaudon, D., Shnider, S. J., Tischfield, D. J., Galazo, M. J. & Macklis, J. D. RORβ induces barrel-like neuronal clusters in the developing neocortex. Cereb. Cortex 22, 996–1006 (2012).

28. Bakken, T. E. et al. Single-nucleus and single-cell transcriptomes compared in matched cortical cell types. PLoS One 13, e0209648 (2018).

29. Tripathi, V. et al. The nuclear-retained noncoding RNA MALAT1 regulates alternative splicing by modulating SR splicing factor phosphorylation. Mol. Cell 39, 925–938 (2010).

30. Tushev, G. et al. Alternative 3’ UTRs Modify the Localization, Regulatory Potential, Stability, and Plasticity of mRNAs in Neuronal Compartments. Neuron 98, 495–511.e6 (2018).

31. Crow, M., Paul, A., Ballouz, S., Huang, Z. J. & Gillis, J. Characterizing the replicability of cell types defined by single cell RNA-sequencing data using MetaNeighbor. Nat. Commun. 9, 884 (2018).

32. Stuart, T. et al. Comprehensive Integration of Single-Cell Data. Cell 177, 1888–1902.e21 (2019).

33. Qiu, X. et al. Single-cell mRNA quantification and differential analysis with Census. Nat. Methods 14, 309–315 (2017).

34. Kiselev, V. Y. et al. SC3: consensus clustering of single-cell RNA-seq data. Nat. Methods 14, 483–486 (2017).

35. Luo, C. & Ecker, J. R. Methyl-C sequencing of single cell nuclei: snmC-seq2 protocol abstract. https://www.protocols.io/view/methyl-c-sequencing-of-single-cell-nuclei-snmc-seq-pjvdkn6 (2018) doi:10.17504/protocols.io.pjvdkn6.

36. Preissl, S., Wang, X. & Ren, B. Sequencing open chromatin of single cell nuclei: snATAC-seq protocol abstract. (2018) doi:10.17504/protocols.io.pjudknw.

37. Luo, C. et al. Single nucleus multi-omics links human cortical cell regulatory genome diversity to disease risk variants. bioRxiv 2019.12.11.873398 (2019) doi:10.1101/2019.12.11.873398.

38. Cao, J. et al. Joint profiling of chromatin accessibility and gene expression in thousands of single cells. Science 361, 1380–1385 (2018).

39. Zhu, C., Preissl, S. & Ren, B. Single-cell multimodal omics: the power of many. Nat. Methods 17, 11–14 (2020).

40. Butler, A., Hoffman, P., Smibert, P., Papalexi, E. & Satija, R. Integrating single-cell transcriptomic data across different conditions, technologies, and species. Nat. Biotechnol. 36, 411–420 (2018).

41. Haghverdi, L., Lun, A. T. L., Morgan, M. D. & Marioni, J. C. Batch effects in single-cell RNA-sequencing data are corrected by matching mutual nearest neighbors. Nat. Biotechnol. 36, 421–427 (2018).

42. Welch, J. D. et al. Single-Cell Multi-omic Integration Compares and Contrasts Features of Brain Cell Identity. Cell 177, 1873–1887.e17 (2019).

43. Lodato, S. et al. Gene co-regulation by Fezf2 selects neurotransmitter identity and connectivity of corticospinal neurons. Nat. Neurosci. 17, 1046–1054 (2014).

44. Xie, W. et al. Epigenomic analysis of multilineage differentiation of human embryonic stem cells. Cell 153, 1134–1148 (2013).

45. Peukert, D., Weber, S., Lumsden, A. & Scholpp, S. Lhx2 and Lhx9 determine neuronal differentiation and compartition in the caudal forebrain by regulating Wnt signaling. PLoS Biol. 9, e1001218 (2011).

46. Hon, G. C. et al. Epigenetic memory at embryonic enhancers identified in DNA methylation maps from adult mouse tissues. Nat. Genet. 45, 1198–1206 (2013).

47. Yin, Y. et al. Impact of cytosine methylation on DNA binding specificities of human transcription factors. Science 356, (2017).

48. Wingender, E., Schoeps, T. & Dönitz, J. TFClass: an expandable hierarchical classification of human transcription factors. Nucleic Acids Res. 41, D165–70 (2013).

49. He, Y. et al. Improved regulatory element prediction based on tissue-specific local epigenomic signatures. Proc. Natl. Acad. Sci. U. S. A. 114, E1633–E1640 (2017).

50. Gray, L. T. et al. Layer-specific chromatin accessibility landscapes reveal regulatory networks in adult mouse visual cortex. Elife 6, (2017).

51. Sorensen, S. A. et al. Correlated gene expression and target specificity demonstrate excitatory projection neuron diversity. Cereb. Cortex 25, 433–449 (2015).

52. Tasic, B. Single cell transcriptomics in neuroscience: cell classification and beyond. Curr. Opin. Neurobiol. 50, 242–249 (2018).

53. Huang, Z. J. & Paul, A. The diversity of GABAergic neurons and neural communication elements. Nat. Rev. Neurosci. 20, 563–572 (2019).

54. Harris, K. D. et al. Classes and continua of hippocampal CA1 inhibitory neurons revealed by single-cell transcriptomics. PLoS Biol. 16, e2006387 (2018).

55. Barkas, N. et al. Wiring together large single-cell RNA-seq sample collections. bioRxiv 460246 (2018) doi:10.1101/460246.

56. Clark, S. J. et al. scNMT-seq enables joint profiling of chromatin accessibility DNA methylation and transcription in single cells. Nat. Commun. 9, 781 (2018).

57. Hu, Y. et al. Simultaneous profiling of transcriptome and DNA methylome from a single cell. Genome Biol. 17, 88 (2016).

58. Harris, J. A. et al. Anatomical characterization of Cre driver mice for neural circuit mapping and manipulation. Front. Neural Circuits 8, 76 (2014).

59. Madisen, L. et al. A robust and high-throughput Cre reporting and characterization system for the whole mouse brain. Nature Neuroscience vol. 13 133–140 (2010).

60. Tasic, B. et al. Adult mouse cortical cell taxonomy revealed by single cell transcriptomics. Nat. Neurosci. 19, 335–346 (2016).

61. Lein, E. S. et al. Genome-wide atlas of gene expression in the adult mouse brain. Nature 445, 168–176 (2007).

62. Fang, R. et al. Fast and Accurate Clustering of Single Cell Epigenomes Reveals Cis-Regulatory Elements in Rare Cell Types. bioRxiv 615179 (2019) doi:10.1101/615179.

63. Dobin, A. et al. STAR: ultrafast universal RNA-seq aligner. Bioinformatics 29, 15–21 (2013).

64. Lawrence, M. et al. Software for computing and annotating genomic ranges. PLoS Comput. Biol. 9, e1003118 (2013).

65. McGinnis, C. S., Murrow, L. M. & Gartner, Z. J. DoubletFinder: Doublet Detection in Single-Cell RNA Sequencing Data Using Artificial Nearest Neighbors. Cell Systems vol. 8 329–337.e4 (2019).

66. Li, H. & Durbin, R. Fast and accurate short read alignment with Burrows-Wheeler transform. Bioinformatics 25, 1754–1760 (2009).

67. Satpathy, A. T. et al. Massively parallel single-cell chromatin landscapes of human immune cell development and intratumoral T cell exhaustion. Nat. Biotechnol. 37, 925–936 (2019).

68. Wolock, S. L., Lopez, R. & Klein, A. M. Scrublet: Computational Identification of Cell Doublets in Single-Cell Transcriptomic Data. Cell Syst 8, 281–291.e9 (2019).

69. Luo, C. et al. Robust single-cell DNA methylome profiling with snmC-seq2. Nat. Commun. 9, 3824 (2018).

70. Krueger, F. & Andrews, S. R. Bismark: a flexible aligner and methylation caller for Bisulfite-Seq applications. Bioinformatics 27, 1571–1572 (2011).

71. Traag, V. A., Waltman, L. & van Eck, N. J. From Louvain to Leiden: guaranteeing well-connected communities. Sci. Rep. 9, 5233 (2019).

72. Blondel, V. D., Guillaume, J.-L., Lambiotte, R. & Lefebvre, E. Fast unfolding of communities in large networks. J. Stat. Mech. 2008, P10008 (2008).

73. McInnes, L., Healy, J. & Melville, J. UMAP: Uniform Manifold Approximation and Projection for Dimension Reduction. arXiv [stat.ML] (2018).

74. Maaten, L. van der & Hinton, G. Visualizing Data using t-SNE. J. Mach. Learn. Res. 9, 2579–2605 (2008).

75. Wolf, F. A., Angerer, P. & Theis, F. J. SCANPY: large-scale single-cell gene expression data analysis. Genome Biol. 19, 15 (2018).

76. van Dijk, D. et al. Recovering Gene Interactions from Single-Cell Data Using Data Diffusion. Cell 174, 716–729.e27 (2018).

77. Schultz, M. D. et al. Human body epigenome maps reveal noncanonical DNA methylation variation. Nature 523, 212–216 (2015).

78. Zhang, Y. et al. Model-based analysis of ChIP-Seq (MACS). Genome Biol. 9, R137 (2008).

79. He, Y. et al. Improved regulatory element prediction based on tissue-specific local epigenomic signatures. Proc. Natl. Acad. Sci. U. S. A. 114, E1633–E1640 (2017).

80. He, Y. et al. Spatiotemporal DNA Methylome Dynamics of the Developing Mammalian Fetus. bioRxiv 166744 (2017) doi:10.1101/166744.

81. Fornes, O., et al. JASPAR 2020: update of the open-access database of transcription factor binding profiles. Nucleic Acids Res. (2019) doi:10.1093/nar/gkz1001.

82. Grant, C. E., Bailey, T. L. & Noble, W. S. FIMO: scanning for occurrences of a given motif. Bioinformatics 27, 1017–1018 (2011).

